# NINscope: a versatile miniscope for multi-region circuit investigations

**DOI:** 10.1101/685909

**Authors:** Andres de Groot, Bastijn J.G. van den Boom, Romano M. van Genderen, Joris Coppens, John van Veldhuijzen, Joop Bos, Hugo Hoedemaker, Mario Negrello, Ingo Willuhn, Chris I. De Zeeuw, Tycho M. Hoogland

## Abstract

Miniaturized fluorescence microscopes (miniscopes) have been instrumental to monitor neural activity during unrestrained behavior and their open-source versions have helped to distribute them at an affordable cost. Generally, the footprint and weight of open-source miniscopes is sacrificed for added functionality. Here, we present NINscope: a light-weight, small footprint, open-source miniscope that incorporates a high-sensitivity image sensor, an inertial measurement unit (IMU), and an LED driver for an external optogenetic probe. We highlight the advantages of NINscope by performing the first simultaneous cellular resolution (dual scope) recordings from cerebellum and cerebral cortex in unrestrained mice, revealing that the activity of both regions generally precedes the onset of behavioral acceleration. We further demonstrate the optogenetic stimulation capabilities of NINscope and show that cerebral cortical activity can be driven strongly by cerebellar stimulation. To validate the performance of our miniscope to image from deep-brain regions, we recorded in the dorsal striatum and, using the IMU to assess turning movements, replicate previous studies that show encoding of action space in this subcortical region. Finally, we combine optogenetic stimulation of distinct cortical regions projecting to the dorsal striatum, to probe functional connectivity. In combination with cross-platform control software, NINscope is a versatile addition to the expanding toolbox of open-source miniscopes and will aid multi-region circuit investigations during unrestrained behavior.

## Introduction

Cellular resolution imaging using miniaturized fluorescence microscopes (miniscopes) permits the monitoring of the topology of activity in brain circuits during unrestrained behaviors. While advances in electrophysiology now enable recordings from many thousands of neurons at once in awake animals (Juavinett et al., 2018; Jun et al., 2017), imaging approaches can sample the activity of individual neurons and retain information about how their activity is spatially distributed in a large network (Kim et al., 2016; Stirman et al., 2016; Terada et al., 2018). Often an anatomical substrate exists for clustered activity such as is the case in the cerebellum, where nearby Purkinje cells receive input from climbing fibers originating in adjacent neurons of the inferior olive brainstem nucleus (Ruigrok, 2010). Thus, imaging approaches can reveal how individual cells embedded in a larger network display coordinated activity during different stages of behavior or training (Galiñanes et al., 2018; Giovannucci et al., 2017; Heffley et al., 2018; Wagner et al., 2017). Moreover, because of their ability to record in freely moving animals, miniscopes have been instrumental in uncovering neural activity patterns occurring during natural behaviors and related brain-states including social interactions (Kingsbury et al., 2019; Liang et al., 2018; Murugan et al., 2017; Remedios et al., 2017) or sleep (Chen et al., 2018; Cox et al., 2016) with fully intact vestibular input.

Open-source miniscopes have provided affordable tools to probe cellular activity in rodents (Cai et al., 2016; Ghosh et al., 2011) and birds (Liberti et al., 2017, 2016) during unrestrained behavior and have so far allowed recordings from a single region. However, in order to understand how brain circuits elicit behavior, it would be preferable to probe multi-region interactions during either spontaneous or trained behaviors. If miniscopes are used to this end, they should be sufficiently light and compact to allow recordings from more than one site without compromising image quality, permit straight-forward behavioral tracking, and have the ability to drive circuits optogenetically. To address these needs, we have developed a versatile and compact miniscope (NINscope) with a sensitive CMOS sensor, integrated inertial measurement unit (IMU) and an accurate LED driver for optogenetic actuation of other brain regions using a custom-made LED probe.

Leveraging the capabilities of NINscope, we demonstrate functional interactions between the cerebellum and cortex in unrestrained mice wearing dual miniscopes. Complex spike activity in Purkinje cell dendrites correlated with cellular activity measured in the cortex during periods of movement, in line with previous anatomical and functional studies of cerebello-thalamo-cortical connectivity (Akkal et al., 2007; Badura et al., 2018; Bostan et al., 2013; Gao et al., 2018; Hoover and Strick, 1999; Wagner et al., 2019). Moreover, activity correlated across regions typically preceded peak behavioral acceleration. Using NINscope’s built-in optogenetic stimulation capabilities in conjunction with accelerometer read-out, we show that activation of the cerebellar hemispheres or vermis elicits lateralized behavioral responses and activation of cortical neurons reiterating cerebello-cerebral functional connectivity.

Finally, we demonstrate the applicability of NINscope to image from neurons in the mouse dorsal striatum, a deep-brain region accessible only after implantation of a relay GRIN lens. Using NINscope’s accelerometer and behavioral analysis of video data, we identify cells in the dorsal striatum whose activity is exclusively modulated when mice make turns contralateral to the recording site, highlighting lateralization in the dorsal striatum and the role of these neurons in representing action space (Barbera et al., 2016; Cui et al., 2013; Klaus et al., 2017). Optogenetic stimulation of prefrontal or secondary motor cortex together with deep-brain imaging further reveals that inputs from both cortical areas can modulate the activity of neurons in the dorsal striatum, albeit with differing efficacy.

Altogether NINscope permits new types of recordings in unrestrained mice. We demonstrate the feasibility to record from two regions in the same mouse concurrently, to use a built-in LED driver in combination with an implantable LED probe for optogenetic stimulation, and to parse behavioral states by the inclusion of an accelerometer. NINscope is an open-source project enabling others to build on its design and functionalities, thereby contributing to a growing range of open-source tools to study neural circuits during unrestrained behavior.

## Results

### NINscope design and functionality

NINscope (**Figure 1A-C**) follows the same general design as other open-source miniscopes, with a number of space and weight saving modifications. A thinner optical emission filter and dichroic mirror (both 500 µm) were used and the emission filter (1 mm) was glued to a plano-convex lens using optical bonding glue (NOA81, Norland Products). Custom high-density interconnector (HDI) sensor and interface PCBs (10 by 10 mm, **Figure 1D**) were designed, fitted with electronic components using a pick-and-place machine (NeoDen4, NeoDen Tech, Hangzhou, China) and stacked to maintain a small footprint. This allowed for inclusion of an IMU and multiple LED drivers including one for optogenetic actuation and two for driving excitation LEDs. The latter feature allows for the extension from single to dual excitation mode.

**Figure 1.**
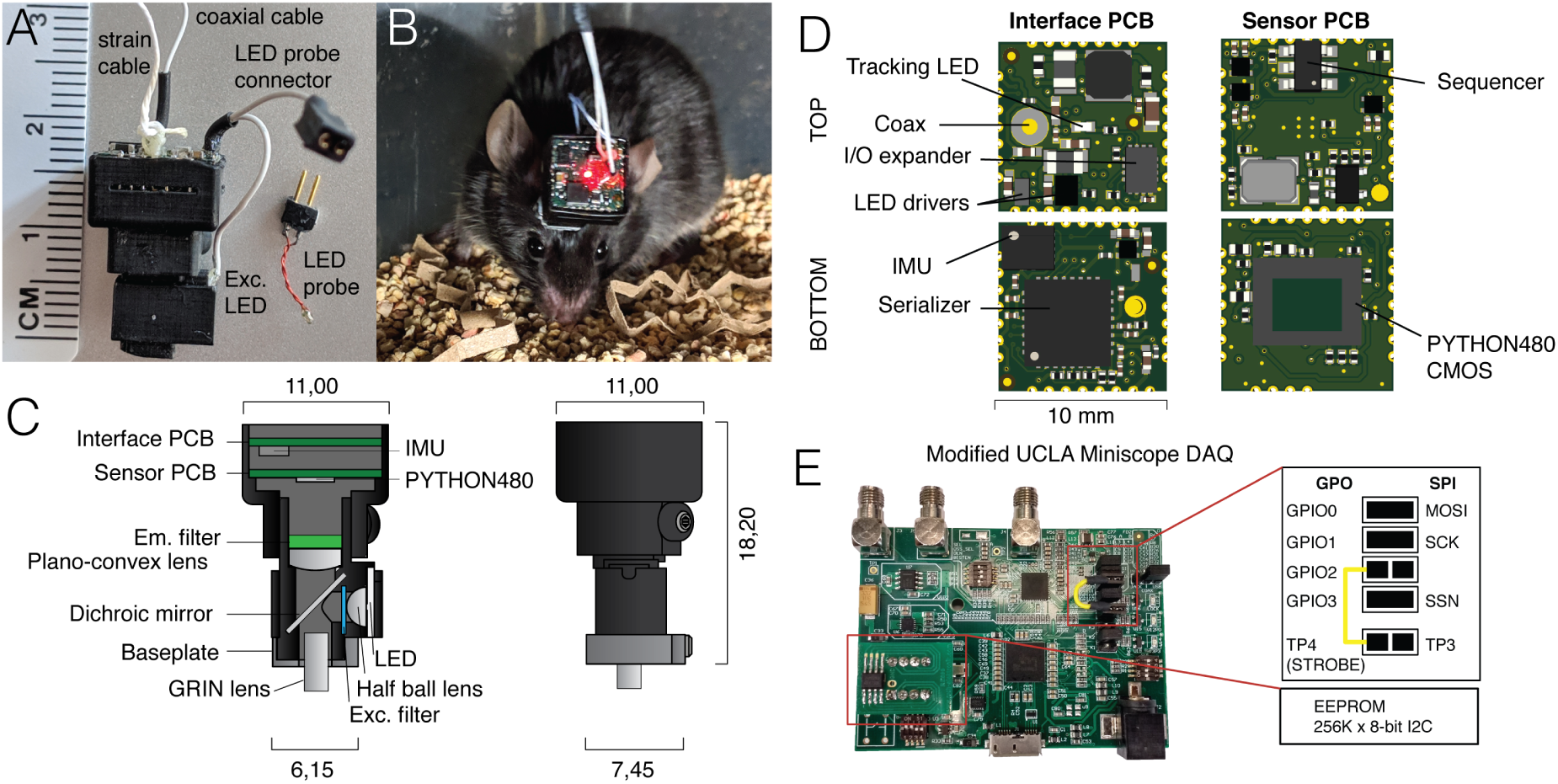
NINscope, a versatile and compact miniscope. **A**, Photograph of NINscope with coaxial and strain cables, excitation (exc.) LED wire, optogenetic LED probe, as well as the probe wire with connector (LED probe connector) coming from the miniscope. **B**, Mouse with a mounted NINscope revealing the red tracking and indicator LED. **C**, Schematic drawings of the NINscope with dimensions in mm. Two 10 by 10 mm HDI PCBs, one for interfacing with the data acquisition box, the other containing the CMOS imaging sensor, are stacked and mounted in a 3D printed enclosure. Excitation light from an LED is collimated with a half ball lens, passes through an excitation filter and is reflected by the dichroic mirror onto the specimen. The emitted fluorescence is collected through the GRIN objective lens, and passes the dichroic and a plano-convex lens, which focuses an image onto the CMOS sensor. An emission (Em.) filter is glued onto the plano-convex lens with optical bonding glue. **D**, KiCad drawings of the custom-built interface and sensor PCBs with top and bottom views including all labeled components. The interface PCB contains an inertial measurement unit (IMU) for measuring head acceleration and orientation, 3 LED drivers including one for optogenetic (strobe) control and 2 for excitation LEDs (1 used), as well as a red tracking LED, the serializer, and the IO expander. The sensor PCB contains the PYTHON480 CMOS sensor and sequencer, which controls sensor timing and the image core. **E**, The UCLA Miniscope DAQ card v3.2 was used with minor modifications including a 256 kB x 8-bit I2C EEPROM (STMicroelectronics) and a wired connection, bridging general purpose input/output 2 (GPIO2) with test point 4 (TP4). The serial peripheral interface (SPI) signals: master output slave input (MOSI), serial clk (SCK) and slave select (SSN) are connected to GPIO0, GPIO1 and GPIO3 through jumpers.

The PYTHON480 sensor (ON Semiconductor) was chosen as a compact yet sensitive CMOS sensor (pixel size: 4.8 µm, dynamic range > 59 dB) with modest power requirements. NINscope uses a 1.8 mm diameter GRIN lens (numerical aperture 0.55, #64-519, Edmund Optics) as objective and has a magnification of ∼4.6×. This gave us approximate field sizes of 786 by 502 μm, using a 752 x 480 pixels Region Of Interest (ROI). In software, we implemented the option to translate this ROI to cover the CMOS sensor area of 800 x 600 pixels (836 by 627 μm).

Because of the widespread accessibility of the first generation UCLA Miniscope, we retained the data acquisition (DAQ V3.2) module of the UCLA Miniscope project with minor modifications that included an EEPROM to store a larger modified version of the latest Cypress EZ-USB FX3 firmware and a wired connection from general purpose input/output 2 (GPIO2) to test point 4 (TP4), allowing 1 ms timing accuracy for the optogenetic LED driver (**Figure 1E**). The firmware of the DAQ module was modified to enable serial control over optogenetic and excitation LED brightness, as well as gain, exposure and black level of the CMOS sensor.

The microscope housing was 3D printed (EnvisionTec Micro Plus Advantage printer, RCP30 M resin and Formlabs Form 2 printer, RS-F2-GPBK-04 black resin) to allow for initial rapid prototyping of various miniscope designs. Printing accuracy proved sufficient for our final design enabling us to keep the weight of the miniscope down to 1.6 grams (housing+ optics+PCBs), while permitting the use of two miniscopes simultaneously on one mouse. The top half of NINscope has an enclosure for the sensor and interface PCBs to protect them from damage during unrestrained animal behavior. NINscope is secured onto a small-footprint baseplate (6.5 by 7.5 mm, **Supplemental Figure 1**) with a set-screw. The lower half of the NINscope housing has a protrusion that fits in a notch in the baseplate for increased stability.

A user interface was written in the Processing language (http://processing.org), allowing cross-platform interoperability and control of experimental recordings. The option was included to record both in single or dual head (two miniscope) mode in combination with an additional USB webcam for video capture of behavior. We used ring buffers in the software to avoid frame time delays and frame drops during acquisitions. Timestamps for acquired frames were logged for post-hoc synchronization. The accelerometer data from the IMU can be visualized live during the miniscope recordings. In addition, optogenetic stimulation patterns and LED probe current can be adjusted through the interface providing integrated control of all aspects of the experiment (**Supplemental Figure 2**). A more detailed description of the hardware and software is provided in the Materials and Methods section. Design files and instructions on hardware assembly, firmware programming and software installation can be found at our GitHub site: https://github.com/ninscope.

In order to validate our miniscope, we imaged under various recording and stimulation configurations, including single and dual scope modes across different brain regions. Behavior of animals was monitored with a USB webcam (**Figure 2A**), the miniscope tracking LED (**Figure 2B**), and an onboard accelerometer. In the example shown, Purkinje cells in lobule V of the cerebellum (AP: −6.4, ML: 0 mm) were transduced with GCaMP6f, and their dendrites were imaged and segmented using the CNMF_E algorithm (Zhou et al., 2018) (**Figure 2 C,D**). Minimal light power (∼120-240 μW before, 30-140 μW after entering the GRIN objective, well within the linear range of the excitation LED driver, **Supplemental Figure 3**) was required to obtain high quality signal-to-noise recordings using a GRIN objective mounted on the brain surface (**Figure 2E, Movie 1**). No signal bleaching or photodamage was observed after our imaging sessions (typically 5.000-20.000 frames, i.e., 2-11 minutes per session repeated with brief intermissions at least 10 times). The x, y and z accelerometer channels were used to detect movement onsets (**Figure 2F**) and discern specific behaviors such as rearing (**Figure 2G**), eating and grooming.

**Figure 2.**
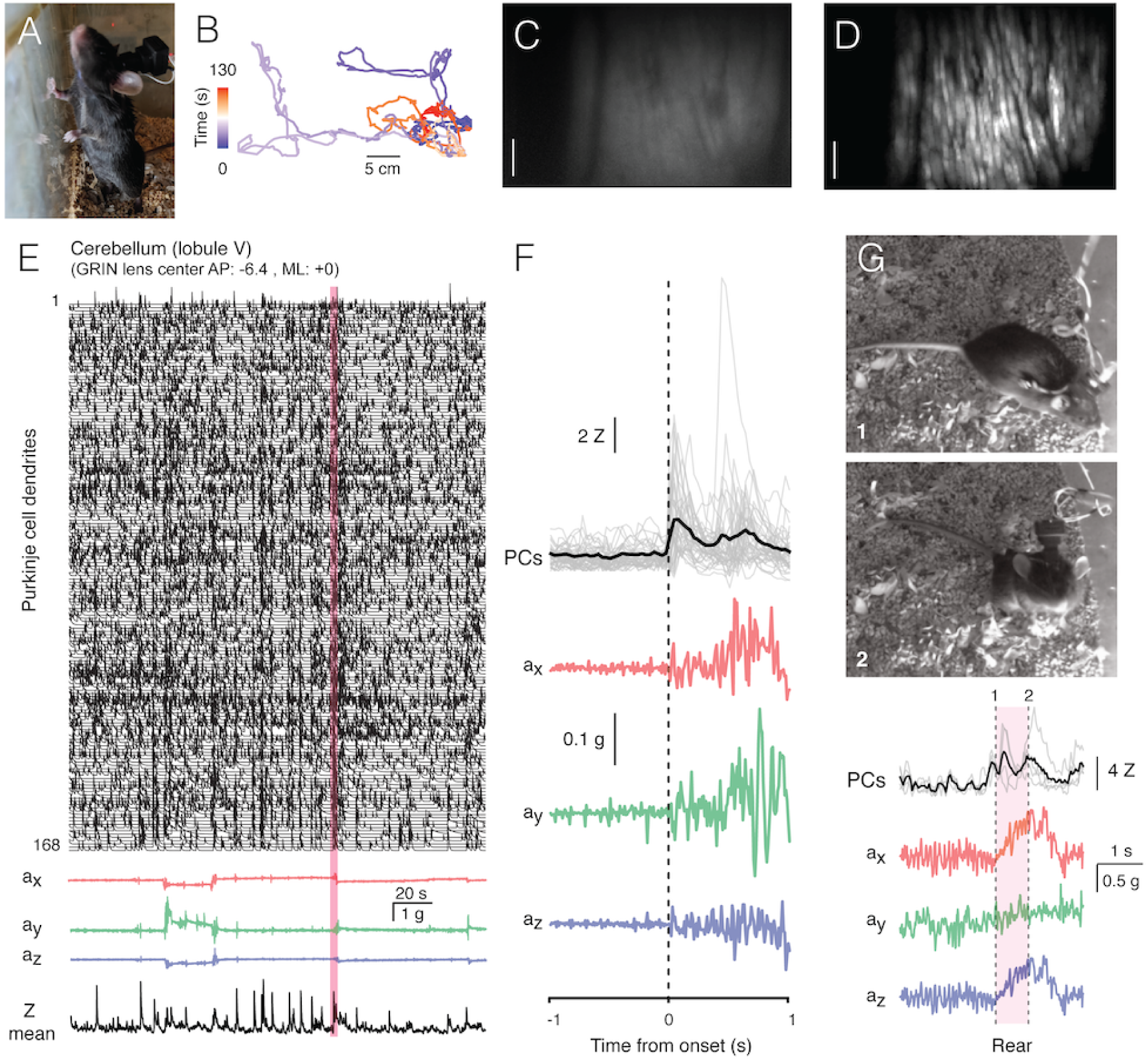
Data acquisition with NINscope. **A**, A mouse wearing a single NINscope mounted over lobule V of cerebellum, which was used to obtain the data shown. **B**, The NINscope tracking LED can be used to obtain information about animal position when combined with concurrent webcam recordings. Colors represent the time from the start of recording and the track represents cage exploration over the course of ∼ 2 minutes. **C**, Average projection of NINscope frames acquired in cerebellum lobule V (scale bar = 100 µm). **D**, Maximum projection of Purkinje cell dendrites that were segmented using CNMF_E (Zhou et al., 2018). **E**, Calcium transients obtained from 168 segmented Purkinje cell dendrites shown together with the x,y and z accelerometer data and the mean Z-scored signal revealing periods of Purkinje cell co-activity. **F**, Purkinje cell transients were aligned to acceleration onset in the red shaded area in E. In this example a reflexive movement (twitch) was triggered by a loud clap. **G**, Spontaneous behaviors monitored with a webcam can be associated with distinct signatures in the accelerometer data such as during rearing. The dotted lines indicate the time points corresponding to the animal just prior (1) and after rearing (2). A subset of Purkinje cells also responded during this behavior.

### Cellular resolution imaging of cerebello-cerebral interactions in unrestrained mice

An important motivation for building lighter and smaller miniscopes is the ability to record from two regions concurrently in unrestrained mice. Our specific goal was to obtain concurrent cellular resolution recordings of cerebellum and cerebral cortex. There is ample anatomical evidence for cerebello-thalamo-cerebral loops (Akkal et al., 2007; Bostan et al., 2013; Hoover and Strick, 1999) and an increasing number of studies suggest that functional interactions within such loops are important for the proper expression of social, cognitive and motor (planning) behaviors (Badura et al., 2018; Gao et al., 2018; Stoodley et al., 2017). The ability to record from both cerebellum and cortex simultaneously in unrestrained mice opens up the possibility to study how these interactions play out during natural spontaneous behaviors.

Given the reduced footprint and weight (1.6 g), we were able to mount two NINscopes on a single mouse. Under these conditions, mice displayed typical cage exploration, grooming, eating and rearing as seen in mice with one miniscope (**Figure 3A**). Our design allows recordings from two regions in mice with a minimum inter-baseplate distance of 8.15 mm and miniscopes placed at an angle of 15° with respect to each other (**Figure 3B**). Given these constraints concurrent recordings from frontal aspects of cortex (encompassing premotor cortex (M1), secondary motor cortex (M2) and prefrontal cortex) and cerebellum, but also from cerebellum and rostral striatum become possible. Thus, with NINscope, one can record from multiple sets of different areas involved in the planning, initiation and control of movement.

**Figure 3.**
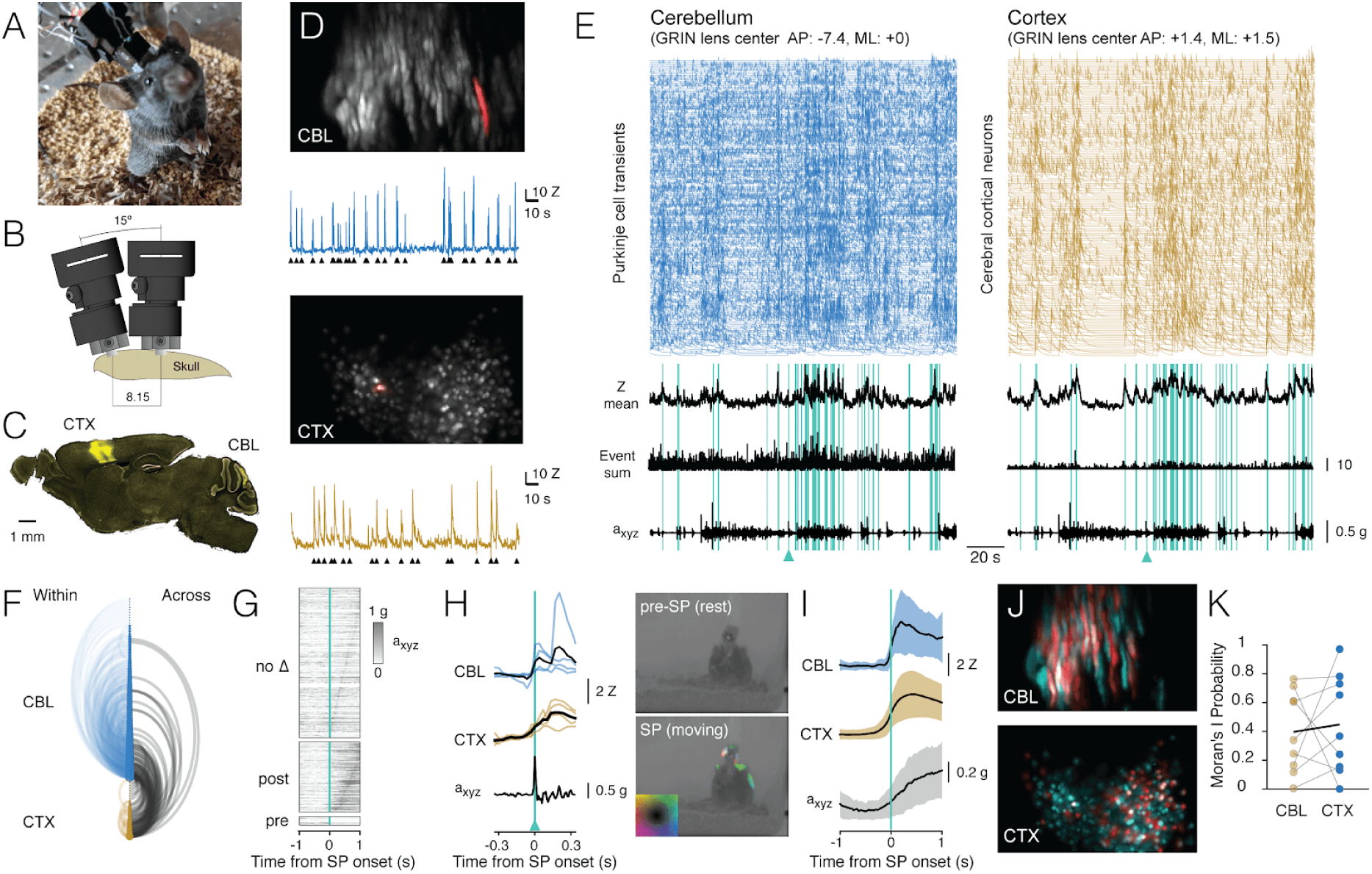
Cellular resolution imaging of cerebello-cerebral interactions in unrestrained mice. **A**, Mouse with two mounted NINscopes. **B**, A minimal inter-baseplate distance of 8.15 mm is achieved when two NINscopes are placed at a 15° angle. **C**, GCaMP6f was transduced selectively in cerebellar Purkinje cells (lobule VI or simplex lobule) and non-selectively in motor cortex. **D**, Examples of segmented Purkinje cell dendrites in the cerebellum (CBL) and neurons in cortex (CTX) with a cell in each area highlighted in red with their corresponding calcium transients and detected events (black triangles). **E**, Dual site recordings from cerebellar lobule VI and motor cortex showing responses of all segmented neurons in each region with the Z-mean scored signal, number of co-active Purkinje cell dendrites and compound acceleration signal (a_xyz_,√(x^2^+y^2^+z^2^)). The cyan lines represent epochs where synchronous patterns (SPs) were found across cerebellum and cortex. **F**, Combined arc plots for the dataset visualizing intra-cerebellar and intra-cerebral (within) and cerebellar-cerebral (across) SP connectivity for the data shown. Node radii scale by the number of cells that a node connects to. In this example it is clear that cerebellar neurons with high within SPs also display significant SPs across regions. **G**, SPs were used to trigger the compound acceleration signal. Behavioral acceleration could be assigned to four categories consisting of no change (no Δ, 64%, sorted by peak response), behavioral acceleration post-SP (31%), pre-SP (4%), or around-SP (1%, not shown). **H**, Across-regions SPs are associated with significant deviations from baseline in the accelerometer compound signal. Responsive cells are shown in cerebellum (lobule VI) and cortex triggered off of the SP. A mouse resting prior to an SP that made a (left, upward) movement around SP onset. Animal movement visualized with optic flow is color-coded (inset shows directional color plane). **I**, Population averaged responses triggered around detected SPs reveal responses in cerebellum and cortex during accelerometer upslope. **J**, Cells in the cerebellum and cortex participating in an SP (red) and cells that did not (cyan). **K**, Little spatial clustering across SPs - as assessed using Moran’s I - was observed across all recordings. Shown are the Moran’s I probabilities calculated for cerebellum and cortex during SPs.

For our dual site recordings virus injections were performed to selectively transduce GCaMP6f in cerebellar Purkinje cells of cerebellar simplex lobule (GRIN objective center AP:-5.8, ML:2.2 mm) or lobule VI (GRIN objective center AP:-7.4, ML:0.0 mm) and globally in neurons of motor cortex (GRIN objective center AP: +1.4, ML: 1.5) covering large parts of the caudal forelimb area and a small part of M2 at the rostral border of the GRIN objective lens (**Figure 3C**).

We were able to extract signals from hundreds of cells in both areas (cerebellum: 141±53, range 62-200, cortex: 201±90, range 89-361, mean± SD, n=4 mice, **Figure 3D**). Calcium transients from Purkinje cell dendrites expressing GCaMP6f displayed faster decay kinetics than cells in cortex expressing the same calcium sensor protein (t_1/2_ cerebellum: 0.217±0.12s, 15784 transients, t_1/2_ cortex: 0.488±0.34s, 9424 transients, mean±SD, Hedge’s G = 1.19, **Figure 3D, E**), likely reflecting differences in endogenous calcium buffering (Baimbridge et al., 1992; Celio, 1990). Event rates measured from Purkinje cell dendrites corresponded to the underlying rate of climbing fiber input (0.62±0.39 Hz, mean±SD, 1629 cells, n=4 mice), while cortical neurons fired events at a lower rate (0.22±0.28 Hz, mean±SD, 2212 cells, n=4 mice).

In all recordings we observed distinct periods of increased activity across regions (**Movie 2)** around periods of animal movement (**Figure 3E**) as gauged from the compound acceleration signal and inspection of the raw data. To quantify correlations across cerebellum (lobule VI, simplex lobule each 2 animals) and cortex, transient event times were concolved with an Epanechnikov kernel and summed over all cells, resulting in a kernel sum, where global synchronous patterns (SPs) were defined as instances where the kernel sum exceeded mean+2σ (cyan lines in **Figure 3E**). A visualization of the functional connectivity of cells is shown in the combined arc plots of **Figure 3F**, where node size scales with the number of SPs within cerebellum (lobule VI in this example), cortex and across the two regions. Both cerebellar Purkinje cells and cortical neurons displayed SPs within and across regions with Purkinje cells having significantly more within-SPs than cortical neurons (fraction of total lobule VI: 66.4±18.9%, simplex lobule: 73.3±27.4%, cortex: 35.7±17%, mean±SD, CBL vs CTX p = 2.7801e-17, KS test). Significantly more Purkinje cells than cortical neurons participated in across-SPs (fraction of total lobule VI: 42.3±29.6%, fraction simplex lobule: 47.4±31.6%, cortex: 26.7±18.5%, mean±SD, CBL vs CTX p = 1.8409e-15, KS test).

We next looked into whether correlated activity in cerebellum and cortex were associated with behavioral acceleration. To determine if it preceded or followed across-region-SPs, the compound acceleration signal was triggered off of across-SPs. We could assign behavioral acceleration to four categories (**Figure 3G**), consisting of no significant change (64%, 206/321 across-SPs), behavioral acceleration post-SP (31%, 99/321), pre-SP (4%, 13/321) or around-SP (1%, 3/321). Thus, the majority (86%, 99/115) of SPs associated with significant acceleration events followed the across-SP. We further confirmed that SPs are associated with acceleration events by comparing the fraction of randomly triggered versus SP-triggered acceleration exceeding mean+2σ of baseline to show that the fraction of SP-triggered acceleration events could not have arisen by chance (random vs SP-triggered, p=0.0017, KS test).

As expected, calcium transients in cerebellar Purkinje cells and cortical neurons occurred around behavioral acceleration (**Figure 3H,I**). In the example shown (triggered off an across-SP indicated by the cyan triangles in **Figure 3E and 3H**), a large deflection occurred in the compound accelerometer signal around an SP onset. This movement was also seen when inspecting the concurrent webcam image frames and visualized here by optic flow, with colors representing direction of movement before (top) and around the SP onset (bottom).

Aligning recordings (n=4 animals, 9 recordings) to post-SP onsets revealed a clear upslope of the accelerometer signal coinciding with calcium increases in both cerebellar Purkinje cells and cortical neurons (**Figure 3I**). Although calcium imaging does not provide very accurate latency estimates, we did examine the relative timing of the calcium responses in cerebellum and cortex relative to movement acceleration. Latencies to peak response of the calcium transients in cerebellum, cortex and the compound acceleration signal revealed that the maximum acceleration came after the peak response in both cerebellum and cortex (CBL: 444±280 ms, CTX: 505±260 ms, a_xyz_: 700±269 ms, Kruskal-Wallis test p=1.6553e-08). A post-hoc test using Sheffe’s S revealed that the mean ranks for the calcium transient peak latencies were both significantly different relative to the delayed peak latency of the accelerometer signal, while this did not apply when comparing the mean ranks of cerebellar and cortical calcium transients. Thus, based on our preliminary data recorded with NINscope, we show that coordinated cerebello-cerebral activity generally precedes peak acceleration of a movement.

In some recordings we observed spatial clustering of activity in cerebellum and/or cortex during across-SPs (**Figure 3J**). To quantify spatial clustering of cells participating in an across-SP, the spatial autocorrelation measure Moran’s I was calculated. The Moran’s I probability, the probability that the highly co-firing cells are randomly distributed, was computed. These probabilities (cerebellum: 0.40±0.28, range = 0.005-0.76, cortex: 0.45± 0.34 range = 0-0.97) suggested that there was no apparent spatial clustering of SP activity (**Figure 3K**). Future studies parsing more precisely annotated behavior using a larger cohort of animals could resolve the occurrence of spatial clustering during behaviorally relevant SPs.

### Cerebral cortical responses during multi-site cerebellar stimulation

So far, miniscope optogenetics has focused on stimulating within the same field-of-view (Stamatakis et al., 2018). Stimulating within the same field-of-view sets constraints on the miniscope design necessitating additional optics, which adds weight and increases the footprint of a miniscope. It also requires the use of an opsin that can be spectrally separated from the activity indicator. Moreover, it is often desirable to stimulate at a site distal to the miniscope, for example to activate indirect projection pathways. By directly driving the LED from the miniscope, the amount of cabling can be reduced, while stimulus onsets can be directly logged to disk together with the image frames and accelerometer data to ease post-hoc analysis. Using the integrated LED driver of NINscope we demonstrate its use by stimulating cerebellar Purkinje cells in transgenic Pcp2-Cre Jdhu x Ai32(RCL-ChR2(H134R)/EYFP) mice while performing calcium imaging from neurons in motor cortex transduced with GCaMP6f (**Figure 4A**, GRIN objective center AP: +1.4, ML: 1.5). We chose 4 cerebellar locations to implant the blue LEDs (470 nm): Crus II ipsi- and contralateral to the imaging site on the right hemisphere, contralateral simplex lobule and lobule VI in medial cerebellar vermis (**Figure 4B**). LED probes consisted of the implanted LED itself, a wire and a connector with pins (**Figure 4C**) that allowed switching the miniscope LED driver connection from one implant site to the next (**Figure 4D**).

**Figure 4.**
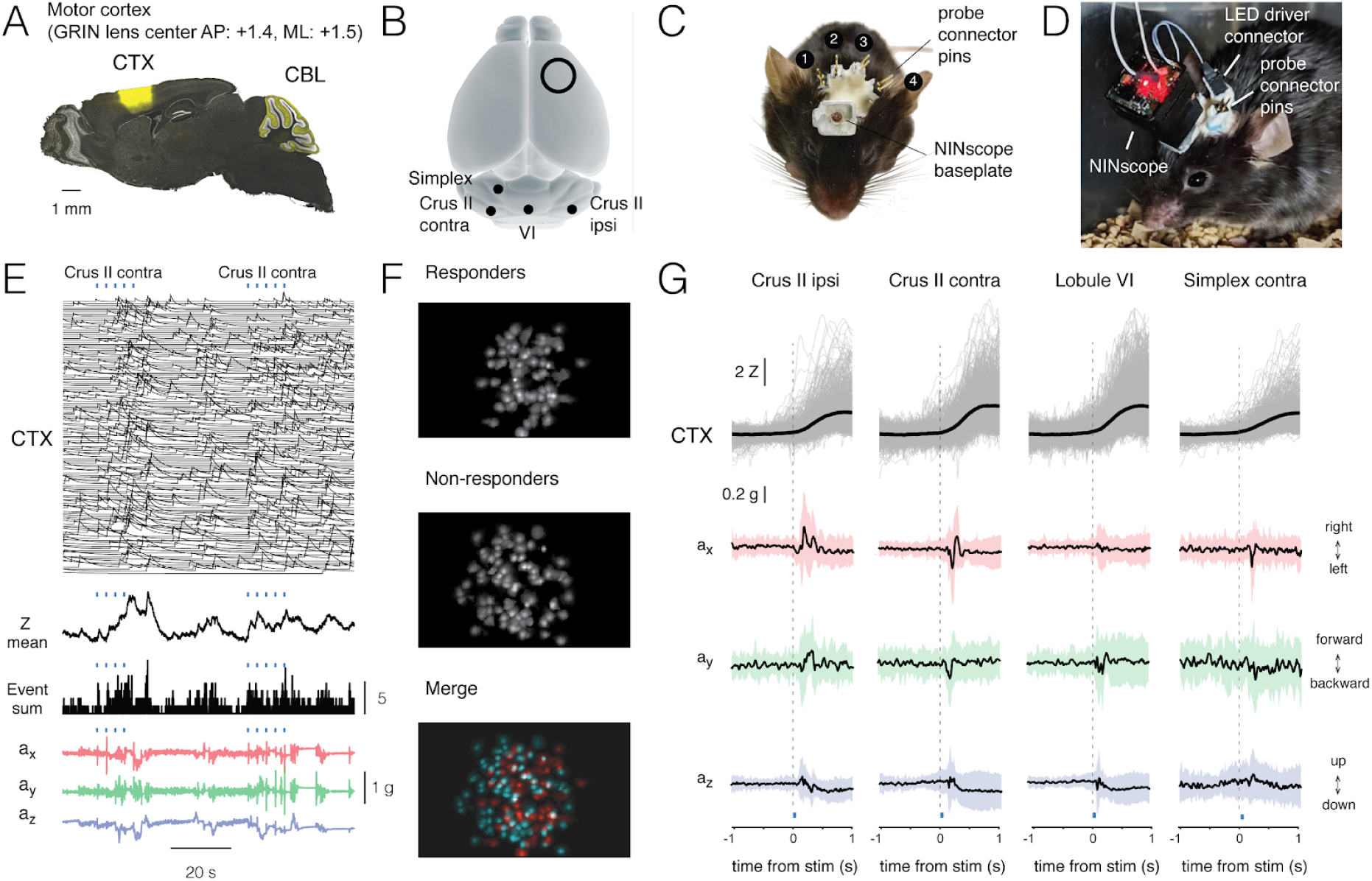
Cerebral cortical responses to multi-site cerebellar stimulation. **A**, Pcp2-Cre Jdhu mice were crossed with Ai32 mice to obtain selective expression of ChR2(H134R) in cerebellar Purkinje cells. Neurons of the motor cortex were transduced in these mice with GCaMP6f. **B**, LEDs (470nm) were implanted at four sites over the cerebellar surface with locations corresponding to simplex lobule, area crus II contralateral to the imaging site, vermis lobule VI and area crus II ipsilateral to the imaging site. **C**, Mouse with a baseplate above the imaging site and the four connector pins that were used to connect the NINscope to the LEDs. **D**, Mouse with a mounted NINscope and connection to one of the stimulation sites. **E**, Optogenetic stimulation of Purkinje cells (50 ms, 22 mA, 2.3 mW) evoked clearly discernible increases in both mean response and number of co-active cells in motor cortex (a subset of calcium transients are shown for clarity). Repeated stimulation induced ramp-like activity in cortex. The accelerometer data revealed deflections following stimulus offset likely reflecting rebound cerebellar nuclear output after release from Purkinje cell inhibition. **F**, In this example a large fraction of cells responded (responders) to optogenetic stimulation of contralateral Crus II. Color merge shows spatial localization of responders (red) and non-responders (cyan). Responsive cells were selected using the criterion that the post-stimulus signal had to exceed mean+2*σ* of the pre-stimulus baseline. **G**, Calcium transients (gray: individual transients, black: mean) triggered to stimulus onset at four different locations over the cerebellar surface and corresponding x, y and z channel accelerometer data. Crus II stimulation reveals lateralization of the behavioral response with stimulation on the right eliciting rightward and stimulation on the left leftward head movements. Evoked behavioral reflexes generally occurred prior to calcium transient onsets in the cerebral cortex.

Stimulation of each of the cerebellar sites implanted with LEDs (50 ms, 22 mA current, 2.3 mW light power) evoked responses in motor cortex of the right hemisphere (**Figure 4E-G, Movie 3**). Recurring cerebellar stimulation (0.3 Hz, 4-5 times) could induce ramp-like activity as seen when averaging responses across cells in motor cortex (**Figure 4E**). The miniscope accelerometer data registered movement that occurred with a delay after stimulus offset (crus II ipsi: 76±23ms; crus II contra: 80±30 ms; lobule VI: 79±34 ms; simplex lobule: 73±32 ms, mean±SD, n=2 mice) in line with earlier observations that disinhibition of Purkinje cells and associated rebound excitation of neurons in the cerebellar nuclei can elicit delayed motor reflexes (Hoebeek et al., 2010; Witter et al., 2013). A significant number of cells met the criterion of displaying calcium increased exceeding mean+2*σ* of the pre-stimulus baseline and were labeled as responders (**Figure 4F**). Across all recordings and sites of stimulation the number of responders was roughly half of all segmented cells in motor cortex (49.9±14.2%, mean±sd, 2227/4614 cells, n=2 mice). Mean amplitudes of Z-scored calcium transients measured in neurons of motor cortex varied depending on the region stimulated (crus II ipsi: 2.1±1.3 Z; crus II contra: 2.9±1.5 Z; lobule VI: 2.9±1.6 Z; simplex lobule: 2.0±1.3 Z; Kruskal-Wallis test p = 1.1239e-70 and post-hoc multiple comparisons using Sheffe’s S revealed significantly different mean ranks for stimulation of contralateral lobule VI and crus II relative to the other groups) (**Figure 4G**), but all displayed comparable onset times (crus II ipsi: 218±120 ms; crus II contra: 221±118 ms; lobule VI: 233±123 ms; simplex lobule: 212±130 ms, mean±sd; Kruskal-Wallis, p=0.1, n.s) relative to stimulus offset. Transient onset times had non normal distributions as assessed with an Anderson-Darling test (A-D test statistic: 5.76, range: 16-577 ms relative to stimulus offset), suggesting that a subpopulation of cortical cells could be responsive during the execution of the optogenetically triggered motor reflexes. On average, however, cerebellar Purkinje cell stimulation caused slow rising cortical responses that could not have directly contributed to movement, but instead could prime motor cortex for upcoming movements. The most dense projections out of cerebellum cross and one would therefore expect strongest activation of the contralateral cortex. A stronger peak amplitude was seen with contralateral cerebellar stimulation, but a significant response was also evoked with ipsilateral cerebellar stimulation. One possible explanation could be the presence of strong commissural projections between the hemispheres (Chovsepian et al., 2017). The comparatively slow sampling associated with calcium imaging may have masked any inter-hemispheric delays. A caveat of these validation experiments might be that we did not stimulate focally by directing light through small diameter optical fibers; yet even with our coarse stimulation clear lateralization of behavioral responses was observed. Stimulation of Purkinje cells over both cerebellar hemispheres evoked opposing head movements as determined from our accelerometer data with leftward movements when stimulating over left and rightward movements when stimulating over right crus II (**Figure 4G**). Crus I and II in rodents are essential for sensorimotor integration in relation to orofacial and whisking behavior (Ju et al., 2019; Romano et al., 2018), but head and neck information provides input to these two areas as well via the external cuneate nucleus (Huang et al., 2013; Quy et al., 2011) and in humans, head movements co-occur with activation of the cerebellar hemispheres (Prudente et al., 2015). Lateral stimulation of left simplex lobule evoked more modest lateral movements as compared to crus II stimulation, while they were mostly absent when stimulating over medial vermis lobule VI where forward/backward movements were relatively more pronounced. Overall, responses of cerebral cortical neurons were robust irrespective of the stimulated cerebellar region, revealing strong functional cerebello-cerebral connectivity. The behavioral reflexes registered with the accelerometer upon Purkinje cell stimulation did not appear directly correlated with cerebral cortical activation, suggesting a more likely scenario of downstream targets in the brainstem underlying these reflexes.

### Action encoding in dorsal striatum

The striatum is a subcortical structure that is inaccessible to imaging without lowering a GRIN relay lens into the site of interest (**Figure 5A**). Due to tissue damage along and below the lens track, longer recovery times are required before imaging can commence (Bocarsly et al., 2015). A significant amount of light is lost through a combination of two GRIN lenses along the optical path (GRIN relay and GRIN objective lens), thereby rendering these experiments more challenging than imaging from cerebellar Purkinje cell dendrites or superficial layers of cerebral cortex. In order to validate NINscope to study the striatum in unrestrained animals and prove its effectiveness for deep-brain imaging, we revisited previous work that has proposed a role of the dorsal striatum (DS) in contraversive movement initiation and action encoding (Cui et al., 2013; Klaus et al., 2017).

**Figure 5.**
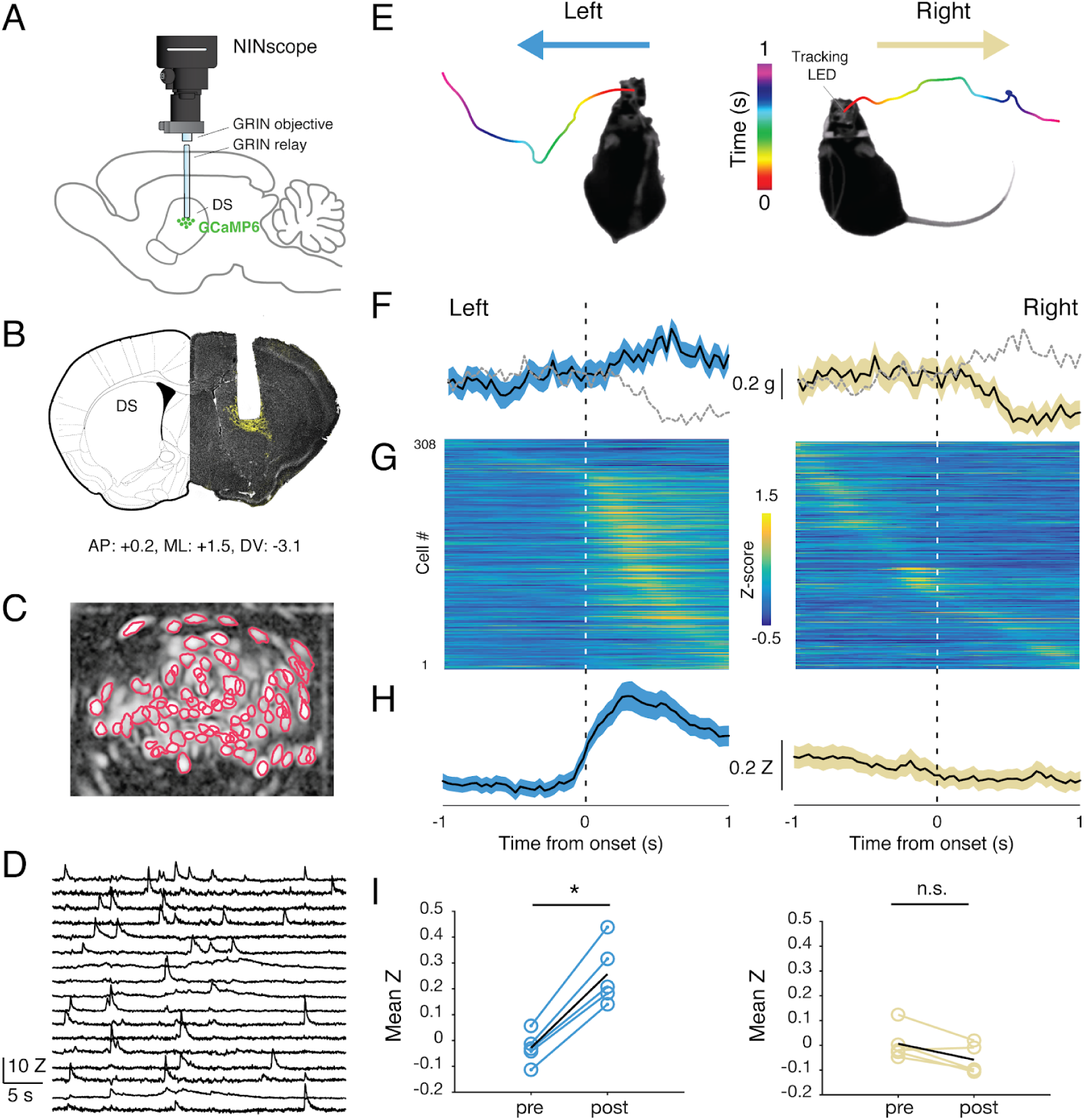
Representation of action space in dorsal striatum. **A**, Schematic of the NINscope configuration, which combines a GRIN objective with a GRIN relay lens to image from the dorsal striatum (DS) of the right hemisphere. **B**, Coronal section showing the GRIN relay lens track and neurons in DS expressing GCaMP6f (yellow). **C**, Segmented neuron ROIs are plotted on top of a correlation image of activity in DS extracted using CNMF-E. **D**, Representative Z-scored calcium transients imaged from DS. **E**, Animals were monitored in an open-field arena using an observational camera. The track-light at the top of the NINscope as well as accelerometer x channel allowed scoring left- and rightward movements. The color-coded lines reflect position of the NINscope tracking LED over time (1 sec). **F**, Accelerometer data showing mean left and right acceleration of the x channel around movement onset (mean±SEM). Dashed gray lines represents contraversive acceleration.. **G**, Sorted Z-scored transients one second before and after movement onset (n=5). A clear increase was seen in neurons of right DS for left turns (contralateral), which was absent during right turns (ipsilateral). **H**, Mean calcium transient responses revealed clear modulation of activity during left turns (mean±SEM). **I**, Quantification of calcium transient responses before and after movement onset for left and right turns, respectively. Right DS only displayed a significant calcium-transient increase for left turns (p<0.05).

Using a viral vector with the human synapsin promoter, we transduced all striatal neurons with GCaMP6 in a caudal, dorsal part of the striatum of the right hemisphere (**Figure 5B**). Directly following viral transduction, a 600 µm diameter GRIN relay lens was implanted above the region of interest. Animals were baseplated after lowering a NINscope with mounted GRIN objective lens and baseplate to just above the GRIN relay lens to bring cells into focus. In DS, we extracted signals from up to 84 cells (62±16.70, mean±sd, range 38-84; **Figure 5C**), which, based on their calcium transients, typically had a rate of around 1 Hz (1.106±0.93 Hz, mean±sd, 308 cells, n=5) (**Figure 5D**). Despite using relatively low light power (∼300 µW after the objective and before the GRIN relay lens), we obtained good signal-to-noise recordings. Using the tracking NINscope tracking LED together with the accelerometer data, we found epochs where mice made both body and head turns (**Figure 5E, Movie 4**). Such turns were associated with up- or downward deflections in the x channel of our accelerometer, reflecting left or right-turning movements, respectively (**Figure 5F**). During left turns (contralateral to imaging location in DS), a majority of neurons in the right DS (85%, 308 cells, n=5) had their peak response after action initiation and 20% displayed significantly elevated responses for the duration of action execution (paired t-test baseline vs action execution activity, p<0.05) (**Figure 5G, H**). The largest of these began after movement initiation, suggesting a predominant association with action execution rather than preparation (latency onset, 310±45 mss, mean±sd). None of the cells we recorded from responded to forward-backward movements or movements ipsilateral to the site of recording (**Figure 5G, H**), confirming lateralization of movement signals in the striatum. When signals were averaged one second pre- and post-movement initiation (Figure 5I, J), repeated measures ANOVA revealed a significant main effect (F_(1.18, 4.7)_=16.48, *p=0*.*01*) of action execution. Post-hoc Bonferroni correction for multiple comparisons demonstrated that the effect occurred exclusively during epochs of contralateral (t=9.45, df=4, *p=0*.*001*), but not ipsilateral movement initiation (t=3.07, df=4, *NS*).

### Top-down control of dorsal striatum

To reveal the impact of cortical inputs on neuronal activity in DS of unrestrained animals, we optogenetically stimulated the orbitofrontal cortex (OFC) or secondary motor cortex (M2) during DS imaging, taking advantage of the capability of our miniscope to directly trigger an LED. The opsin ChrimsonR was transduced in either of the two cortical regions and a 625 nm LED implanted above the cortex (**Figure 6A**). In reduced preparations, OFC and M2 have both been found to regulate the activity of neurons in specific DS regions (Corbit et al., 2019), with the impact of OFC on DS being stronger than that of M2. We sought to confirm these findings in freely behaving mice. To this end, we first assessed the direct terminal fields of OFC and M2 to DS and mapped this onto a representative brain atlas image (**Figure 6B**). M2 input to DS was more diffuse than the projections of OFC, consistent with previous findings (Corbit et al., 2019; Hintiryan et al., 2016; Hunnicutt et al., 2016). During imaging sessions, animals were able to freely explore an open-field arena while OFC or M2 were optogenetically stimulated (10s, 20Hz frequency, 5µs pulse width, 3.4mW LED). Three distinct response types of DS neurons were observed during OFC and M2 stimulation periods and consisted of decreased, increased, or unchanged (no change) responses, relative to baseline activity (paired t-test baseline vs stimulus evoked activity, p<0.05) (**Figure 6C**). Calcium transients during stimulation differed between these clusters in firing frequency (decrease: 0.23±0.25 Hz, mean±sd, 33 cells, n=4, unchanged: 0.77±0.68 Hz, mean±sd, 199 cells, n=4, increased: 1.63±1.23 Hz, mean±sd, 22 cells, n=4).

**Figure 6.**
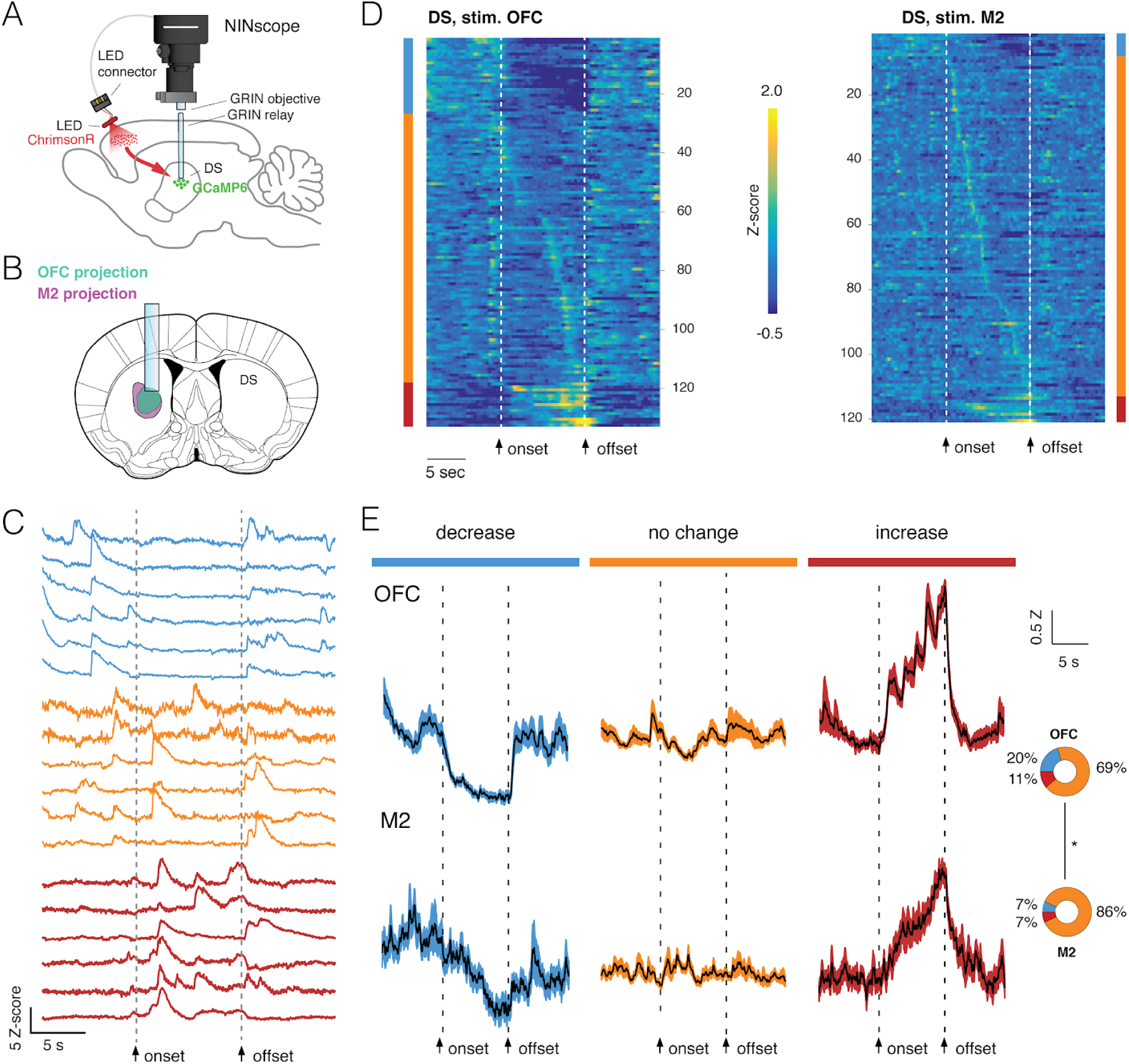
Top-down, optogenetic control of activity in dorsal striatum. **A**, Schematic showing placement of the NINscope with GRIN objective and GRIN relay lens to record from dorsal striatum (DS) as well as location of the optogenetic LED probe driven by the integrated LED driver. Viral vectors were injected either in orbitofrontal cortex (OFC) or secondary motor cortex (M2) to transduce neurons with ChrimsonR, or in DS to transduce neurons with GCaMP6 for calcium imaging. **B**, Terminal fields of OFC and M2 mapped onto a representative brain atlas show overlapping inputs in DS underneath the GRIN relay lens. **C**, Different types of responses were found in DS when either OFC or M2 are optogenetically stimulated. Neurons exhibited decreases of activity (blue), no apparent change (orange), or increased responses (red). **D**, Z-scored calcium transients during OFC (left, n=2) and M2 (right, n=2) stimulation (10 s pulse, 20 Hz). For each neuron, 10 trials were averaged. **E**, Responses averaged over all DS neurons for all 10 trials revealed similar types of modulation during stimulation of OFC and M2 (mean±SEM). The circular inset denotes the fraction of cells that showed suppression, no change, or increased responses.

Stimulation was repeated for 10 trials and responses were averaged over trials (**Figure 6D**). Modulation of activity in subpopulations of DS neurons during OFC stimulation was observed: 20% of the neurons (26/133) displayed a significant decrease in activity, 69% displayed no change (93/133) and 11% (15/133) were increased. M2 stimulation had a comparable, but weaker impact on activity of subpopulations of DS neurons with 7% of neurons (7/107) showing a decrease, 86% (93/107) displaying no change and 7% (7/107) of cells showed an activity increase. Although the fraction of responsive DS neurons (i.e. those showing a decrease or increase) during cortical stimulation significantly differed between OFC (31%) and M2 (14%) (chi-squared(1)=13.85, *p<0*.*001*), the mean response for each category of DS neurons during stimulation was comparable (**Figure 6E**). Decreased neurons showed a gradual reduction in activity for the duration of stimulation, whereas neurons responding to stimulation with increased activity showed a progressive increase as long as the stimulation was provided, suggesting that input modulation of activity in DS neurons scaled with stimulation duration.

## Discussion

### NINscope

We have demonstrated the applicability of NINscope to perform dual site recordings in mice, and combine superficial and deep brain imaging with optogenetic stimulation, while parsing movements through a built-in accelerometer.

Our 1.6 g miniscope does not compromise in terms of weight or footprint to add functionality, which makes it a valuable contribution to the expanding tool chest of open-source miniscopes. Since NINscope control software is platform-independent it can be used on all major operating systems and using different hardware configurations including laptops that are within reach of most users.

Several design choices had to be made during NINscope prototyping. We 3D printed the microscope housing to keep the design light, but eschewed the use of an electrowetting lens, which would have made focusing practical, but the miniscope too heavy and bulky for dual site recordings. The inclusion of a high resolution LED driver for optogenetic stimulation together with the use of an implantable LED probe provided the ability to directly drive projections at their site of origin instead of their terminations within the imaging field-of-view, which is a feature that has not been integrated into a miniscope before.

Given that our building plans are open-source, the core functionality of our scope can be expanded depending on the specific research questions that need to be addressed, which ultimately dictate the size, weight and functionality constraints. The ability to repurpose the UCLA Miniscope DAQ box (v3.2) for use with NINscope underscores the benefits of sharing resources and we hope that our contributions will further advance miniscope development as well.

### Cerebello-cerebral interactions

To our knowledge we present the first concurrent cellular resolution recordings of cerebellum and cerebral cortex in unrestrained mice. We show that synchronous patterns of activity across cerebellum and cerebral cortex are common across multiple behavioral states and are associated significantly with movement acceleration. The ability to record from two sites (e.g. encompassing cerebellum and cortex, or cerebellum and rostral striatum) in freely behaving animals opens up new avenues for research into cerebello-cerebral and cerebello-striatal interactions during the expression of innate and learned behaviors and can help to resolve outstanding questions with regard to the role of the cortical and subcortical structures in planning, learning and executing skilled movements (Gao et al., 2018; Guo et al., 2015; Kawai et al., 2015; Sauerbrei et al., 2018). Changes in joint cell participation during procedural learning (Galiñanes et al., 2018; Guo et al., 2015) can be quantified concurrently in cortical and subcortical structures to determine if cell responses are enhanced or suppressed (Kostadinov et al., 2019). Although we did not find a consistent spatial clustering of activity in cerebellum and cortex when neurons in both regions displayed joint activity, a more careful parcellation of the behavior in a larger sample may reveal that such clustering does indeed occur. In line with findings of tightly coordinated activity in cortex and subcortical structures, recent work has highlighted the motor cortex as an input-driven dynamical system (Sauerbrei et al., 2018), where thalamic input is a prerequisite to drive movements as they happen. Cerebellar output appears essential not only for contributing to the precision of ongoing movements (Becker and Person, 2019), but also their planning in sensory discrimination tasks (Chabrol et al., 2019; Gao et al., 2018). Thus, despite redundancy of motor systems as seen when lesioning selective regions- and assessment of motor performance after a recovery period-, both cortical and subcortical regions are required for successful motor planning and performance under natural conditions. Using NINscope we performed a crude assessment of functional connectivity between cerebellum and cerebral cortex by optogenetically stimulating Purkinje cells bilaterally and medially over the cerebellum. Such stimuli evoked reflexive movements with delays similar to those that we have reported previously and correspond with the timing of rebound activity in the cerebellar nuclei after stimulus offset (Witter et al., 2013). Here, we found that the site of cerebellar stimulation dictates the direction of a movement as shown by our accelerometer data. This movement directionality coincides with the known anatomy of the cerebellum where each cerebellar hemisphere controls the ipsilateral side of the body. We further found that about half of the neurons from cortical imaging sessions responded significantly to strong synchronous cerebellar activation with onsets that were delayed relative to the stimulus onset. Thus, these responses seem to signal the occurrence of a movement rather than that they can be associated with their execution.

### Action encoding and input driven responses in dorsal striatum

The PYTHON480 CMOS sensor incorporated in NINscope enabled recording signals from the dorsal striatum (DS) with good signal-to-noise at powers of a few hundred µW before the relay GRIN lens confirming the applicability our miniscope to image from deep-brain structures. By analyzing the data collected both via video camera and the built-in accelerometer, we found that a large fraction of recorded neurons in the right DS responded exclusively during leftward turning (contralateral) behavior of the mice, confirming previous findings that identified the striatum as a structure representing action space (Barbera et al., 2016; Cui et al., 2013; Klaus et al., 2017; Tecuapetla et al., 2014). Although our results are overall consistent with previous work, we did not find evidence for spatial clustering of responsive neurons when mice made contralateral turns.

The DS receives dense input from the ipsilateral orbitofrontal (OFC) and secondary motor (M2) cortices. Slice physiology experiments, using comparable neuroanatomical coordinates as in this study, have proposed that synaptic inputs from OFC onto neurons of DS have a stronger impact on the excitability of medium spiny neurons than M2 inputs (Corbit et al., 2019). Using NINscope in combination with optogenetic stimulation of OFC or M2 at their site of origin, we now confirm this finding in vivo showing that, also in the intact circuit, more DS neurons respond to OFC than M2 stimulation. Both inputs modulated activity in DS neurons in awake behaving mice, but the fraction of responsive neurons was larger for stimulation of the OFC projection. Here, we did not disambiguate the types of neurons that were positively or negatively modulated. Increases of activity in DS neurons are potentially driven by feedforward excitation directly onto striatal medium spiny neurons (Hintiryan et al., 2016) while direct input onto fast-spiking interneurons (Mallet et al., 2005) could underlie decreased responses in DS neurons during our stimulation. Irrespective of the sign of modulation, we found responses that scaled with the duration of the stimulus, thus, proving the efficacy of both stimulation and imaging.

In conclusion, NINscope is a versatile open-source miniscope that could fit a niche of users that desire dual region recordings in unrestrained animals, OS interoperability, optogenetic manipulation of areas at a distance from the image site, behavioral parsing using an accelerometer, or a combination of all of the above.

## Materials and Methods

### Printed circuit board design

NINscope uses two printed circuit boards (PCBs): a CMOS image sensor and interface PCB each 10 by 10 mm (thickness 0.6 mm, HDI standard) and developed using the open-source and cross-platform electronics design automation suite KiCad (http://kicad-pcb.org). The interface PCB includes the DS90UR913A (Cypress Semiconductor) serializer to connect via an FPD-link III to the DAQ board. The interface PCB is placed on top of the sensor PCB and the two PCBs are connected by wires soldered to the castellated holes at the PCB edges. The sensor PCB contains voltage regulators, a power sequencer and an oscillator necessary for initializing and operating the image sensor. The voltage regulators are ultra-low noise LDO regulators (NCP163, On Semiconductor) and are specced for use in camera applications. The image sensor needs 3 power sources that are provided in a sequence, which we achieve by using a LM3880 power sequencer (Texas Instruments). A 66.6667 MHz low power CMOS clock oscillator (Kyocera Electronics) is used to run the image sensor. In addition to the serializer, the interface PCB has an IMU sensor (LSM6DSLTR, STMicroelectronics), 2 LED drivers (single LED for optogenetic stimuli: LM36011; dual LED for 1 fluorescence excitation LED: LM3643, Texas Instruments) and an I2C I/O expander (FXL6408UMX, ON Semiconductor). The I/O expander controls signals that are not time critical for the image sensor and the tracking/indicator LED on top of the interface PCB. The LED driver and accelerometer are described in more detail below.

### CMOS image sensor

The sensor PCB includes the PYTHON480 (ON Semiconductor) CMOS SVGA image sensor. The PYTHON480 CMOS image sensor is flexible and has many possible configurations. The challenge was to configure it to our demands. Once we figured out how to start the sensor in CMOS instead of LVDS mode we were able to provide the data signals to the general programmable interface (GPIF) of the Cypress USB controller in order to transfer the image data. To optimize power consumption and limit overheating we opted to switch off the Phase-Locked Loop (PLL) to reduce current consumption by 30%. Without the PLL, the CMOS parallel clock output is 4× times lower constraining the amount of pixel data that can be read-out for a given frame rate. We set acquisition rate in the current NINscope to 30 frames per second and the frame size to 752 x 480 pixels to allow read-out of both pixel data and three accelerometer channels from the IMU. Improved thermal dissipation in future iterations of NINscope will make data at higher frame rates up to 120 Hz accessible and allow additional read-out of the gyroscope data from our IMU.

### Data acquisition hardware

The UCLA Miniscope project has the largest open-source miniscope community with a user base of hundreds of users. We made use of the existing data acquisition (DAQ version 3.2) hardware from the UCLA Miniscope project (http://www.miniscope.org) and introduced minor modifications consisting of a 256 kB x 8-bit I2C EEPROM (STMicroelectronics) to hold larger firmware and connections on the DAQ PCB between GPIO2 to TP4 and SPI signals to GPO0-GPO3. The PYTHON480 CMOS sensor connects via an FPD-Link III and coaxial cable to the Cypress USB controller on the DAQ board. The image sensor has an SPI instead of an I2C interface for sensor configuration. The general purpose outputs (GPO) are used since the FPD-link in the UCLA DAQ supports an I2C and not an SPI bus. This requires routing signals from MOSI, SCKand SS to GPO0, GPO1 and GPO3 allowing full control of the image sensor from the firmware of the USB-controller. Since our firmware is larger than the existing EEPROM on the UCLA DAQ board, we upgraded the EEPROM to 256 kB. The NINscope firmware is based on example firmware derived from the Cypress application note AN75779 (modification date: 30/10/2017). The CMOS image sensor which is registered as a USB imaging device is expanded to function as a composite USB device to allow the addition of a virtual serial port. The virtual serial port is used for communication and configuration of the CMOS sensor and LED drivers (e.g. to set image sensor gain, brightness, black levels, LED settings, optogenetic stimulus parameters) as well as for logging of the accelerometer data.

A low-power (0.65 mA in high-performance mode) 3D accelerometer and gyroscope iNemo inertial module (LSM6DSLTR, STMicroelectronics), set of LED drivers (single LED for optogenetic stimuli: LM36011; dual LED for one fluorescence excitation LED: LM3643, Texas Instruments) and I/O expander (FXL6408, ON Semiconductor) are controlled over the I2C bus. The remaining GPO is used to control pulse generation and precise timing of the optogenetic stimuli by adding a wire connection between GPO2 and TP4 in the UCLA DAQ (**Figure 1E**).

The IMU accelerometer has a range of +/-2 g where each bit corresponds to 0.061 mg. A sample rate of 104 Hz is set, which fills a FIFO buffer with x,y,z data that is read out at the end of every image frame and transmitted over the virtual serial connection with the computer.

### LED drivers and LEDs

A single-LED flash driver (LM36011, Texas Instruments) and dual-LED driver (LM3643, Texas Instruments) were used to respectively control the generation of optogenetic pulses in a 470 nm LED probe (LED 150040BS73240, 402 case size, Wurth Electronics) or 630 nm LED probe (APHHS1005SURCK, Kingbright) and a 470 nm fluorescence excitation LED (Excitation LED LXML-PB02 470nm, Lumileds). The optogenetic LED can be adjusted in increments of 11.725 mA and the excitation LED in 1.4mA increments. The maximum current of either LED is dependent on the total current consumption of the camera and type/length of the cable used. A 621 nm LED (SML-P11UTT86R, Rohm) was integrated on top of the interface PCB to allow camera-assisted tracking of animal position and for notification of miniscope connectivity.

### NINscope housing

NINscope housing prototypes were designed using Inventor (AutoDesk) with initial prototypes being printed using a Micro Plus Advantage printer (EnvisionTec) using the RCP30 M resin to allow printing of fine detail. The thinnest functional wall of the final prototype had a thickness of 500 µm. The Form 2 (Formlabs) printer was subsequently used in printing microscope housing with black resin (RS-F2-GPBK-04). The microscope consists of three parts, an upper part to hold the PCBs, plano-convex lens and emission filter, a lower part for housing the optics including the LED die, half ball lens, excitation filter and dichroic mirror and a sliding cover to secure the LED and protect the optical filters. The lower part of the microscope has a small protrusion that locks in a notch of the custom metal baseplate (**Supplemental Figure 1**) and is secured by a set screw to minimize microscope movement. After printing and cleaning the housing with isopropyl alcohol and sand-dusting, the in- and outside housing printed with the EnvisionTec printer was airbrushed (Infinity CR Plus, Harder & Steenbeck, Germany) with a thin coat of black paint (H12 Flat Black, Gunze, Japan).

### Optics

We used custom-diced excitation (ET470/40x 3.5 × 3.5 × 0.5mm, Chroma), dichroic (T495lpxr 3.5 x 5 x 0.5mm, Chroma) and emission filters (ET525/50m, 4 × 4 × 1mm, Chroma), a N-BK7 half ball lens (3 mm diameter, 47-269, Edmund Optics) and a plano-convex lens (4.0mm diameter, 10.0mm focal length, 45-429, Edmund Optics) to focus onto the CMOS image sensor. The emission filter was bonded to this lens with optical adhesive (NOA81, Norland Products). The GRIN lens (NA 0.55, 64-519, Edmund Optics) was implanted for superficial imaging, or glued into the baseplate for deep-brain imaging, and the miniscope was mounted on a baseplate that was cemented to the skull.

### NINscope cabling

To connect the NIN scope to the DAQ box, a thin (0.101 mm) coaxial wire (38awg A9438W-10-nd, Alpha Wire) of 50 cm length was attached to the miniscope on one end and connected through a connector set (ED90265-ND and ED8250-ND connectors, Digi-Key) to a thicker (1.17mm FEP mm) coaxial wire (VMTX Mini,Pro Power) of 2 m at the other end. The 2 m wire was soldered to a 150 mm RG174 coaxial cable assembly with an SMA straight plug (2096227, LPRS) that could be directly attached to the DAQ board. The excitation LED driver was connected to the LED using 22 mm long ultra-thin wire (UT3607,Habia). A 25 mm long wire of the same type was connected to the optogenetic stimulus LED driver on the interface PCB. On one end a connector was attached (851-43-050-10-001000,Mill-Max of ED90265-ND, Digi-Key) to connect to an optogenetic probe.

### NINscope acquisition software

Acquisition software was written in the cross-platform Processing language (https://processing.org/). The software supports acquisition from up to two miniscopes and one USB webcam. The NINscope is controlled via a serial communication port using the Processing serial library, which supports custom microcontroller devices.

The capture module in the video library for Processing version 2.0 was modified by changing YUV to a raw format to take advantage of the full 8-bit scale. To distinguish cameras and NINscopes of the same type a prefix is added to the string list of all attached devices.

G4P, a Processing library (http://www.lagers.org.uk/g4p/) was used to create the software user interface, allowing access to the most common controls to change microscope settings and update parameters such as light intensity, gain, black level, optogenetic stimulus duration, or G-sensor logging.

Within the Processing sketch a capture event function transfers each captured image into a ring buffer for rapid processing of the image and to avoid long hard drive (HD) access times which could cause frame drops. A thread is used to save all images from ring buffer to HD. The size of the ring buffer can be modified in the sketch but is set to 60. The Last 4 pixels of a frame are reserved for a frame counter to monitor dropped frames. Every frame is saved as grayscale tiff image with the smallest possible header to reduce file size.

PCB and mechanical designs, firmware and acquisition software can be found on GitHub at: https://github.com/ninscope

### Mice

We used both male and female C57/B6 mice (weight range: 22-28 gram). In experiments where Purkinje cells were optogenetically stimulated we used two transgenic mice from a cross between an Pcp2-Cre Jdhu (JAX #010536) and the Ai32 mouse line (JAX # 012569) on a C57/B6 background. All performed experiments were licensed by the Dutch Competent Authority and approved by the local Animal Welfare Body, following the European guidelines for the care and use of laboratory animals Directive 2010/63/EU.

### GRIN objective lens implants and virus injections

Prior to surgeries mice were anesthetized with 3% Isoflurane before being transferred to a stereotactic apparatus after which anesthesia was maintained at 1.5% Isoflurane (flow rate: 0.3 ml/min O_2_). For imaging of the cerebral and cerebellar cortex, a GRIN objective lens (1.8 mm diameter, 0.25 pitch, 64-519, Edmund Optics) was implanted on the brain surface. A small incision was made in the skin after shaving hair removal and disinfection of the skin with iodine solution (5%) and alcohol (70%). Lidocaine (100 mg/ml, Astra Zeneca, UK) was then applied to the exposed skull and the periosteum removed. The center coordinates for GRIN lens placement were located and a small ink dot was placed at the correct location relative to bregma (cerebellar Simplex lobule, AP: −5.8 mm ML: 2.2 mm; lobule VI, AP: −7.4 mm, ML: 0.0 mm; cortex, AP: 1.4 mm, ML: 1.5 mm). Coordinates were scaled relative to the mean bregma-lambda distance (of 4.21 mm) as specified in Paxinos mouse brain atlas. Prior to drilling of the bone, mice received i.p. Injections of 15% D-Mannitol in saline (0.55 ml/25gr) to aid diffusion of virus particles after virus injection. A 2 mm circular craniotomy was then drilled centered around the marked location. In between drilling the skull was kept moist with sterile saline. The skull flap and dura were then removed and virus (Cerebellum: AAV1.CAG.FLEX.GCaMP6f / AAV1.CMV.PI.Cre.rBG mixed 1:1 and diluted in saline 1:3; Cortex: AAV1.Syn.GCaMP6f.WPRE.SV40 diluted in saline 1:3, UPenn Vector Core) was injected at four locations. At each location 25 nl of virus was injected once at 350, twice at 300 and once at 250 µm depth at a rate of 25 nl/min. The craniotomy was covered with gelfoam (Pfizer, USA) soaked in sterile saline (0.9 % NaCl, B. Braun Medical Inc., USA). The GRIN lens was lowered using a vacuum holder placed in the stereotactic apparatus until the lens surface touched the brain and then lowered an additional 50 µm. The edges of the craniotomy were sealed with Kwik-Sil (WPI, USA). Dental cement (Super-Bond C&B, Sun Medical, Japan) was then applied around the lens to secure it in place. Kwik-Cast (WPI, USA) was used to cover and protect the lens. At the end of the surgery animals received an s.c. injection of 5mg/kg Metacam.

To access deep-brain region dorsal striatum, GRIN relay lenses with a diameter of 0.6 mm (length: 7.3mm, numerical aperture: 0.45, Inscopix, CA) were implanted (AP: 0.2, ML: 1.5, DV: −3.1). To ease implantation and reduce damage to tissue above dorsal striatum, a track was created using a 25G needle. Using a motorized stereotaxic arm, the needle was lowered slowly over the course of 10 min, left in place for 10 min, and then retracted with the same speed at a constant rate of 0.31 mm/min. Virus was delivered by lowering a Hamilton syringe over the course of 4 minutes to inject 500 nl of AAVdj.hSyn.GCaMP6s or AAV1.hSyn.GCaMP6f into the dorsal striatum at a rate of 100 nl/min after which the syringe was left in place to increase diffusion of virus (5 min) and subsequently retracted (5 min). Subsequently, the GRIN relay lens was lowered (0.10 mm/min) into the dorsal striatum (DV: −3.1). The gap between the GRIN relay lens and the skull was covered with cyanoacrylate glue and the lens was secured to the skull using dental cement. The GRIN relay lens was covered and protected using Kwik-Cast (World Precision Instruments, USA). A 1.8 mm GRIN objective lens (64-519, Edmund Optics, UK) was secured onto the miniscope baseplate using cyanoacrylate glue and the miniscope was then mounted on the baseplate. Mice were head restricted using a custom-built running belt to allow active locomotion, the scope with baseplate and objective GRIN lens mounted was then lowered above the GRIN relay lens until cells in the field of view became visible, the baseplate was secured to the skull with dental cement (coated with black nail polish) and the miniscope removed.

### LED implants above orbitofrontal and secondary motor cortex

In experiments imaging from dorsal striatum, AAV5.hSyn.ChrimsonR.tdTomato was injected in either the orbitofrontal cortex (AP: 2.8, ML: 1, DV: −2.2) or secondary motor cortex (AP: 2.3, ML: 0.7 DV: −1.2). An LED probe, coated with biocompatible epoxy, designed to connect to NINscope with a red LED (645nm, KingBright Inc., USA), was placed on the brain surface and the LED was secured to the skull using dental cement.

### LED implants above cerebellum

For cerebellar stimulation, a small 1 mm craniotomy was drilled above the cerebellar region of interest (Simplex lobule, AP: −5.8 mm, ML: 2.2 mm; Lobule VI, AP: −7.4 mm, ML: 0.0 mm; Crus II, AP: −7.3 mm, ML: + and −2.7 mm;) following similar sterile procedures as described for the GRIN objective lens implant. The LED -covered in a layer of biocompatible epoxy-was inserted with proper orientation in the craniotomy, which was then sealed with a layer of Kwik-Sil (WPI, USA) followed by a layer of Kwik-Cast. The remaining probe wire and the probe connector were subsequently cemented to the skull with dental cement (Super-Bond C&B, Sun Medical, Japan).

### Imaging

Recordings from cerebellum and cerebral cortex were performed in the mouse home cage, while those obtained from dorsal striatum occurred in a custom open field arena (30 by 30 by 45 cm). The miniscopes were attached either following brief anesthesia (3% isoflurane induction, 30 min. post-anesthesia recovery time) to allow connection of the optogenetics LED driver connector to the LED probe, mounting of one or two scopes and focal adjustment, or -when using a modified baseplate or custom head bar-in a head-fixed condition where animals could run on a treadmill.

### Histology

Mice were transcardially perfused with 4% PFA in 1x PBS (10 mM PO4^3−^, 137 mM NaCl, 2.7 mM KCl). After fixation, brains were removed and kept overnight on 10% sucrose in PBS after which they were subsequently embedded in gelatin and left overnight in 30% sucrose in PBS. Brain sections (50 µm) were cut in a cryostat (LEICA CM3050 S), stained with DAPI and then mounted (Dako Fluorescence Mounting Medium, S3023). Confocal images were collected on a SP8 confocal microscope (Leica, Germany) and whole brain images were obtained by stitching multiple (1024×1024 pixel) acquisitions into a final image.

### Calcium imaging analysis

Timestamps for all scopes were logged to disk and in case of dual scope recordings, frames of the movies were aligned prior to motion correction using NoRMCorre (Pnevmatikakis and Giovannucci, 2017) and signal extraction using CNMF-E (Zhou et al., 2018). In the cerebellar stimulation experiments, neurons in cortex were classified as responders if the post-stimulus signal rose above the pre-stimulus mean+2*σ*. Onset times of these calcium transients were determined by fitting a sigmoid function to the transients and onset time was set to where the fitted function rose above mean+*σ* of the pre-stimulus baseline. For the dorsal striatum experiments, neurons were classified as responders if the signal during the stimulation period differed significantly from the pre-stimulus baseline signal. For every cell, activity pre-stimulus and during stimulation was averaged per trial and statistically tested using paired t-tests. Fiji (Schindelin et al., 2012) was used for raw data inspection and to create movies. Analyses were performed in Matlab (Mathworks, Nantucket), Python 3.7 (van Rossum, 1995) and R (R Core Team, 2019).

### Within and across-region synchronous pattern (SP) detection

Calcium transient events were inferred using a finite rate of innovation algorithm for fast and accurate spike detection (Oñativia et al., 2013) setting *τ*=1 to obtain the onset times of calcium transients at a sub-recording rate resolution from the calcium transients per cell. These onset times were then convolved with an Epanechnikov kernel (steepness = 0.1) and summed over all cells, resulting in a kernel sum. All time intervals for which the kernel sum was two standard deviations above the mean were considered significant global synchronous events. If the onset times of two cells fell into the same synchronous event, these cells were considered to fire synchronously once. For all pairs of cells, we counted how many times these cells both fired inside the same synchronous event. Subsequently, we selected all synchronous pairs that fired together at least five times. These pairs were stored in a graph, where each node represents a cell and each edge the number of shared synchronous firing events. These graphs were converted to an arc diagram using the arcdiagram package (Gaston Sanchez, https://github.com/gastonstat/arcdiagram) in R. Cells were grouped by brain region (cerebellum, cortex) and sorted within the group by graph degree, i.e. the number of cells they correlate to. The degree of correlation is represented by the node radius.

### Analysis of behavioral acceleration

To distinguish whether accelerometer signals exceeded a mean+2σ threshold before or after an SP, we determined first where the largest mean signal occurred. We then made a selection for signals pre- or post-SP that rose above threshold. For the remaining signals we searched 300 ms around the SP (−150, +150 ms) for a rise above threshold. All other signals were classified as showing no SP-related change in behavioral acceleration.

## Author contributions and acknowledgements

AG, JB, JC, JV and TH planned and designed NINscope prototypes. AG developed the electronics and wrote the acquisition software. JB and JV designed and printed mechanical parts. JC simulated and made optical designs. BB, CZ, IW, MN and TH designed experiments. HH and BB performed surgeries and histology. BB and TH performed experiments. BB, RG and TH analyzed the data and/or developed new analysis tools. BB and TH made the figures. AG, BB and TH wrote the paper with input from all authors. TH supervised the project. We thank the GENIE Program at the Janelia Research Campus for generously making their genetically encoded calcium indicators available. We thank the UCLA Miniscope team for making their hardware publicly available, Daniel Aharoni (UCLA) for feedback and discussions. We are grateful to Pieter Roelfsema (NIN-KNAW), Mike Vink (NIN-KNAW) and Kees van Oers (NIOO-KNAW) for their various contributions as part of the NINscope project and Christian Lohmann (NIN-KNAW) for his comments on the manuscript.

## Funding

Financial support for IW was provided by H2020 European Research Council (ERC-2014-STG 638013), the Netherlands Organization for Scientific Research (NWO-ALW, 864.14.010, 2015/06367/ALW). Financial support for CZ was provided by the Netherlands Organization for Scientific Research (NWO-ALW), the Dutch Organization for Medical Sciences (ZonMW), European Research Council (ERC-adv and ERC-POC) and the Royal Netherlands Academy of Arts and Science (KNAW). Financial support for TH was provided by the Royal Netherlands Academy of Arts and Sciences (KNAW) and Top Sector Life Sciences and Health (LSHM18001).

---

**Supplemental Figure 1.**
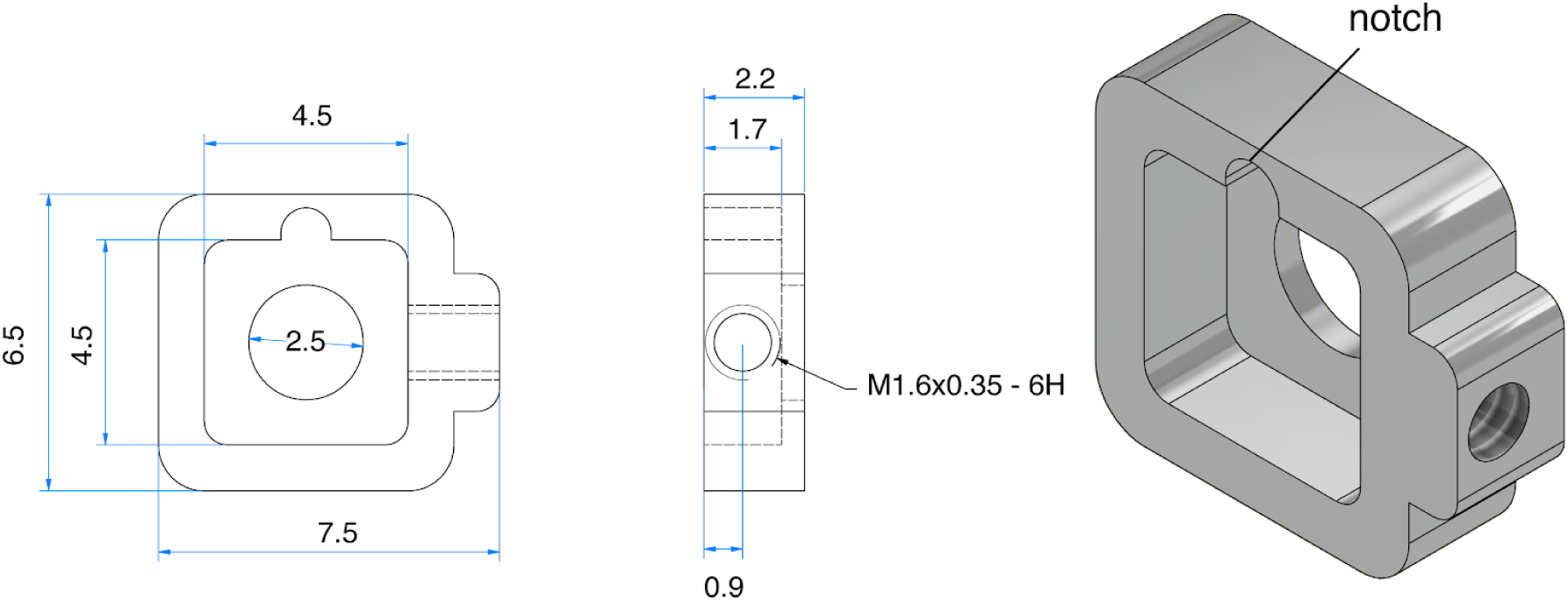
NINscope baseplate. NINscope is secured in a custom designed baseplate using a hexagonal set screw (M1.6×3, 0.7 Hex, part # 913000100016003, Jeveka). The lower half of the NINscope housing has a protrusion that fits in the baseplate notch for added stability.

**Supplemental Figure 2.**
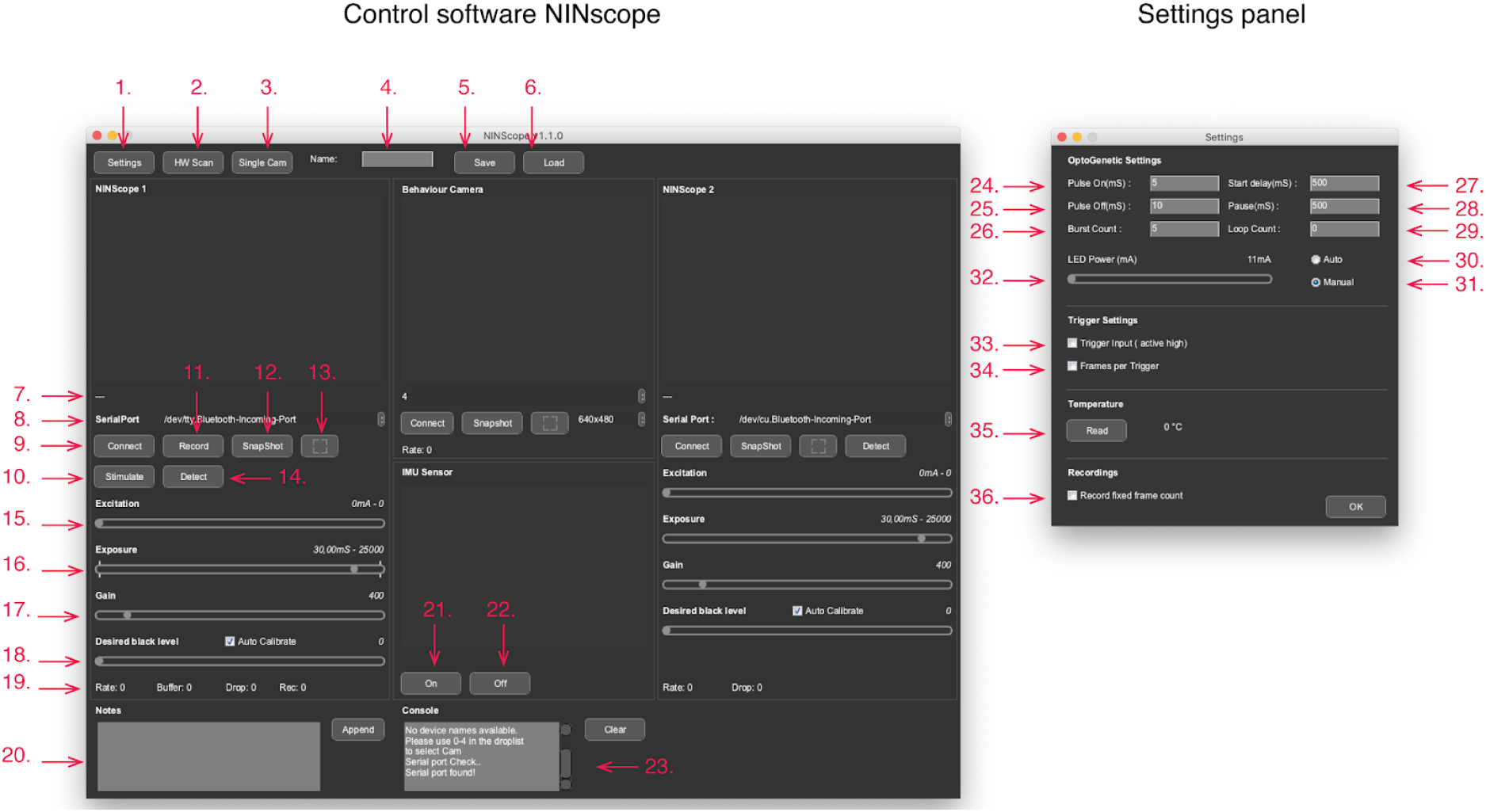
NINscope software. Control software: 1. Access settings panel for optogenetic stimulus parameters, input trigger settings 2. Update list of connected devices (UVC cameras and serial ports) 3. Switch to dual or single cam mode 3. Name of saved settings file 5. Save: save settings file 6. Load: load previously stored settings file 7. Display connected device 8. Serial port selection for device connection 9. Connect or disconnect from a device 10. Initiate manual optogenetic stimulus if enabled 11. Start and stop a recording (disabled in input trigger mode) 12. Take a snapshot 13. Expand camera view 14. Detect which miniscope is connected (tracking LED blinks 5x) 15. Excitation LED current 16. CMOS sensor exposure 17. CMOS gain setting 18. Black level setting (default: auto-calibration) 19. Displays frame rate, number of buffer frames, frame drops and number of frames recorded 20. Text field to add notes, written to disk after pressing the append button. 21. Turn G-sensor on (real-time display in window) 22. Turn G-sensor off 23. Console displays hardware checks upon startup and after a HW scan (see 2.) Settings panel: 24. Optogenetic pulse on duration 25. Optogenetic pulse off duration 26. Optogenetic burst count 27. Optogenetic pulse start delay 28. Optogenetic pulse pause time 29. Optogenetic pulse loop count 30. Start optogenetic stimulation automatically upon record 31. Start optogenetic stimulation upon manual trigger (see 10.) 32. Set current to optogenetic LED 32. Turn on/off trigger input (active high 3.3 V) 33. Set number of frames to record for a trigger input (if not set hold voltage at 3.3 V to keep recording) 34. Read CMOS sensor temperature 35. Record a fixed number of frames for each recording (overrules setting 33). For more information on software usage, please visit https://github.com/ninscope/Software/wiki

**Supplemental Figure 3.**
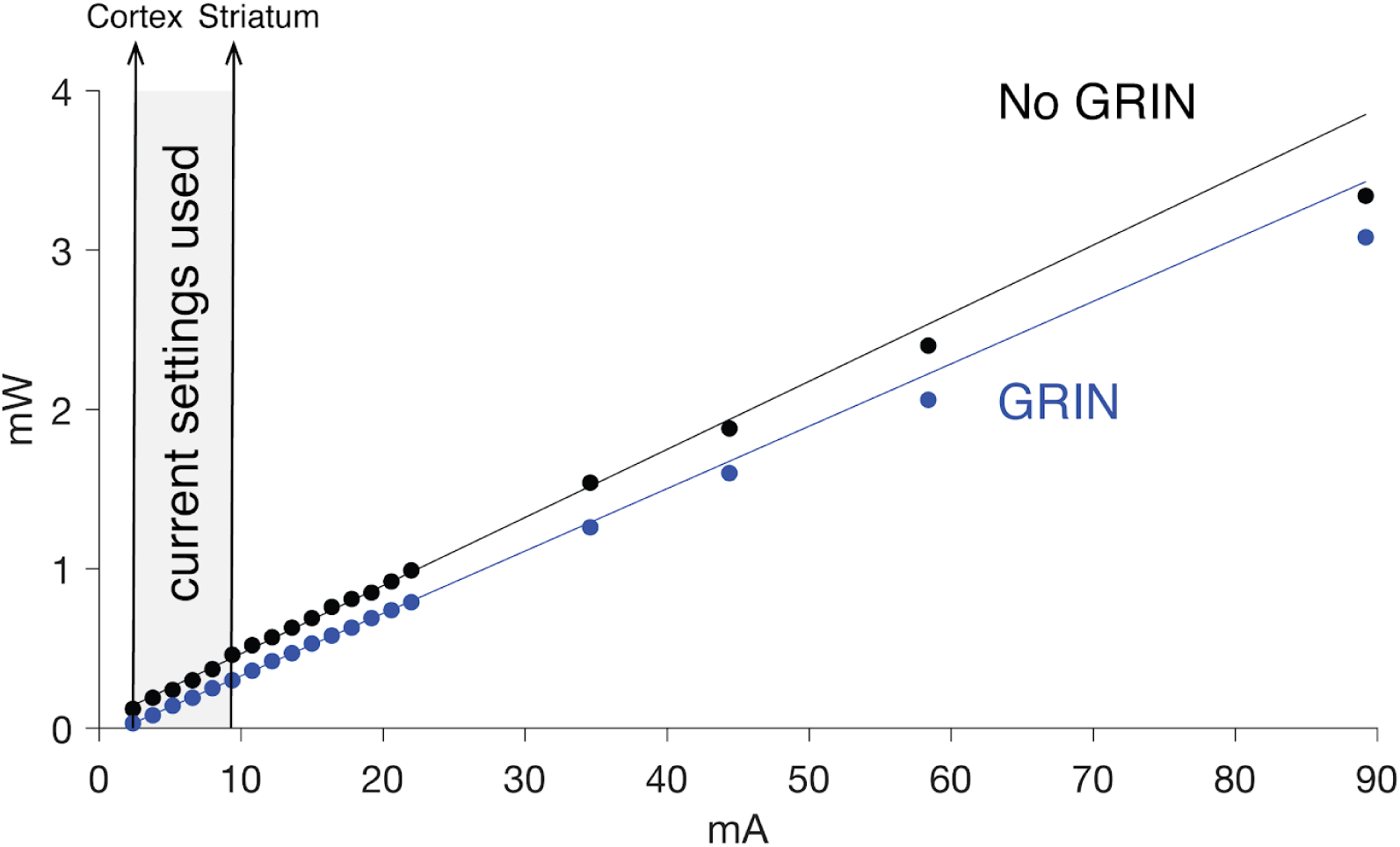
Linearity of excitation LED driver. Excitation LED light power measured as a function of supplied current without a GRIN objective (No GRIN), or after the GRIN objective (GRIN). Arrows indicate the typical excitation LED current settings used to image from cortex or striatum. For striatal imaging light passes through the GRIN objective and an additional GRIN relay lens. We did not measure light transmission through the relay lens.

**Movie 1, Imaging complex spikes with NINscope**. Movie showing activity in Purkinje cell dendrites of cerebellar lobule V measured with NINscope (top) during exploration of the home cage which was monitored with a USB webcam (bottom). Acquisition rate 30 Hz, movie speed 2x.

**Movie 2, Concurrent recordings from cerebellum and cortex using NINscope**. Movie showing activity measured with two NInscopes mounted above motor cortex (top) and cerebellar lobule VI (bottom) during unrestrained behavior. The behavior of the animal was monitored with the accelerometer and a USB webcam. On the right subset of cells (50) from cerebellum (blue) and cortex (ocre) are plotted over the course of the recording. Acquisition rate 30 Hz, movie speed 5x.

**Movie 3, Cerebellar Purkinje cell stimulation and concurrent cortical read-out using NINscope**. Movie showing raw (left) and background subtracted data (right) with the occurrence of the cerebellar stimulus (here 50 ms stimulation of contralateral crus2) indicated by a blue bar. Acquisition rate 30 Hz, movie speed 2x.

**Movie 4, Action encoding in the striatum measured with NINscope**. Activity recorded from dorsal striatum (top) and open-field animal exploration. Acquisition rate 30 Hz, movie speed 1x.

## Notes

#### Summary of Updates

Figure 1 has been updated with a missing component (tracking LED) We have increased the sample size for Figure 6. This is reflected both in the manuscript and Figure. We have made various other corrections and improvements in the Figures, text and added line numbers. Finally, we have included a funding section.

https://github.com/ninscope/

https://scope.nin.knaw.nl

## References

Akkal D, Dum RP, Strick PL. 2007. Supplementary Motor Area and Presupplementary Motor Area: Targets of Basal Ganglia and Cerebellar Output. J Neurosci 27: 10659–10673.

Badura A, Verpeut JL, Metzger JW, Pereira TD, Pisano TJ, Deverett B, Bakshinskaya DE, Wang SS-H. 2018. Normal cognitive and social development require posterior cerebellar activity. Elife 7. doi: 10.7554/eLife.36401

Baimbridge KG, Celio MR, Rogers JH. 1992. Calcium-binding proteins in the nervous system. Trends Neurosci 15: 303–308.

Barbera G, Liang B, Zhang L, Gerfen CR, Culurciello E, Chen R, Li Y, Lin D-T. 2016. Spatially Compact Neural Clusters in the Dorsal Striatum Encode Locomotion Relevant Information. Neuron 92: 202–213.

Becker MI, Person AL. 2019. Cerebellar Control of Reach Kinematics for Endpoint Precision. Neuron. doi: 10.1016/j.neuron.2019.05.007

Bocarsly ME, Jiang W-C, Wang C, Dudman JT, Ji N, Aponte Y. 2015. Minimally invasive microendoscopy system for in vivo functional imaging of deep nuclei in the mouse brain. Biomed Opt Express 6: 4546–4556.

Bostan AC, Dum RP, Strick PL. 2013. Cerebellar networks with the cerebral cortex and basal ganglia. Trends Cogn Sci 17: 241–254.

Cai DJ, Aharoni D, Shuman T, Shobe J, Biane J, Song W, Wei B, Veshkini M, La-Vu M, Lou J, Flores SE, Kim I, Sano Y, Zhou M, Baumgaertel K, Lavi A, Kamata M, Tuszynski M, Mayford M, Golshani P, Silva AJ. 2016. A shared neural ensemble links distinct contextual memories encoded close in time. Nature 534: 115–118.

Celio MR. 1990. Calbindin D-28k and parvalbumin in the rat nervous system. Neuroscience 35: 375–475.

Chabrol FP, Blot A, Mrsic-Flogel TD. 2019. Cerebellar Contribution to Preparatory Activity in Motor Neocortex. Neuron. doi: 10.1016/j.neuron.2019.05.022

Chen K-S, Xu M, Zhang Z, Chang W-C, Gaj T, Schaffer DV, Dan Y. 2018. A Hypothalamic Switch for REM and Non-REM Sleep. Neuron 97: 1168–1176.e4.

Chovsepian A, Empl L, Correa D, Bareyre FM. 2017. Heterotopic Transcallosal Projections Are Present throughout the Mouse Cortex. Front Cell Neurosci 11: 36.

Corbit VL, Manning EE, Gittis AH, Ahmari SE. 2019. Strengthened Inputs from Secondary Motor Cortex to Striatum in a Mouse Model of Compulsive Behavior. J Neurosci 39: 2965–2975.

Cox J, Pinto L, Dan Y. 2016. Calcium imaging of sleep-wake related neuronal activity in the dorsal pons. Nat Commun 7: 10763.

Cui G, Jun SB, Jin X, Pham MD, Vogel SS, Lovinger DM, Costa RM. 2013. Concurrent activation of striatal direct and indirect pathways during action initiation. Nature 494: 238–242.

Galiñanes GL, Bonardi C, Huber D. 2018. Directional Reaching for Water as a Cortex-Dependent Behavioral Framework for Mice. Cell Rep 22: 2767–2783.

Gao Z, Davis C, Thomas AM, Economo MN, Abrego AM, Svoboda K, De Zeeuw CI, Li N. 2018. A cortico-cerebellar loop for motor planning. Nature 563: 113–116.

Ghosh KK, Burns LD, Cocker ED, Nimmerjahn A, Ziv Y, El Gamal A, Schnitzer MJ. 2011. Miniaturized integration of a fluorescence microscope. Nat Methods 8: 871–878.

Giovannucci A, Badura A, Deverett B, Najafi F, Pereira TD, Gao Z, Ozden I, Kloth AD, Pnevmatikakis E, Paninski L, De Zeeuw CI, Medina JF, Wang SS-H. 2017. Cerebellar granule cells acquire a widespread predictive feedback signal during motor learning. Nat Neurosci 20: 727–734.

Guo J-Z, Graves AR, Guo WW, Zheng J, Lee A, Rodríguez-González J, Li N, Macklin JJ, Phillips JW, Mensh BD, Branson K, Hantman AW. 2015. Cortex commands the performance of skilled movement. Elife 4: e10774.

Heffley W, Song EY, Xu Z, Taylor BN, Hughes MA, McKinney A, Joshua M, Hull C. 2018. Coordinated cerebellar climbing fiber activity signals learned sensorimotor predictions. Nat Neurosci 21: 1431–1441.

Hintiryan H, Foster NN, Bowman I, Bay M, Song MY, Gou L, Yamashita S, Bienkowski MS, Zingg B, Zhu M, Yang XW, Shih JC, Toga AW, Dong H-W. 2016. The mouse cortico-striatal projectome. Nat Neurosci 19: 1100–1114.

Hoebeek FE, Witter L, Ruigrok TJH, De Zeeuw CI. 2010. Differential olivo-cerebellar cortical control of rebound activity in the cerebellar nuclei. Proc Natl Acad Sci U S A 107: 8410–8415.

Hoover JE, Strick PL. 1999. The organization of cerebellar and basal ganglia outputs to primary motor cortex as revealed by retrograde transneuronal transport of herpes simplex virus type 1. J Neurosci 19: 1446–1463.

Huang C-C, Sugino K, Shima Y, Guo C, Bai S, Mensh BD, Nelson SB, Hantman AW. 2013. Convergence of pontine and proprioceptive streams onto multimodal cerebellar granule cells. Elife 2: e00400.

Hunnicutt BJ, Jongbloets BC, Birdsong WT, Gertz KJ, Zhong H, Mao T. 2016. A comprehensive excitatory input map of the striatum reveals novel functional organization. Elife 5. doi: 10.7554/eLife.19103

Juavinett AL, Bekheet G, Churchland AK. 2018. Chronically-implanted Neuropixels probes enable high yield recordings in freely moving mice. bioRxiv. doi: 10.1101/406074

Ju C, Bosman LWJ, Hoogland TM, Velauthapillai A, Murugesan P, Warnaar P, van Genderen RM, Negrello M, De Zeeuw CI. 2019. Neurons of the inferior olive respond to broad classes of sensory input while subject to homeostatic control. J Physiol. doi: 10.1113/JP277413

Jun JJ, Steinmetz NA, Siegle JH, Denman DJ, Bauza M, Barbarits B, Lee AK, Anastassiou CA, Andrei A, Aydin Ç, Barbic M, Blanche TJ, Bonin V, Couto J, Dutta B, Gratiy SL, Gutnisky DA, Häusser M, Karsh B, Ledochowitsch P, Lopez CM, Mitelut C, Musa S, Okun M, Pachitariu M, Putzeys J, Rich PD, Rossant C, Sun W-L, Svoboda K, Carandini M, Harris KD, Koch C, O’Keefe J, Harris TD. 2017. Fully integrated silicon probes for high-density recording of neural activity. Nature 551: 232–236.

Kawai R, Markman T, Poddar R, Ko R, Fantana AL, Dhawale AK, Kampff AR, Ölveczky BP. 2015. Motor cortex is required for learning but not for executing a motor skill. Neuron 86: 800–812.

Kim TH, Zhang Y, Lecoq J, Jung JC, Li J, Zeng H, Niell CM, Schnitzer MJ. 2016. Long-Term Optical Access to an Estimated One Million Neurons in the Live Mouse Cortex. Cell Rep 17: 3385–3394.

Kingsbury L, Huang S, Wang J, Gu K, Golshani P, Wu YE, Hong W. 2019. Correlated Neural Activity and Encoding of Behavior across Brains of Socially Interacting Animals. Cell. doi: 10.1016/j.cell.2019.05.022

Klaus A, Martins GJ, Paixao VB, Zhou P, Paninski L, Costa RM. 2017. The Spatiotemporal Organization of the Striatum Encodes Action Space. Neuron 95: 1171–1180.e7.

Kostadinov D, Beau M, Pozo MB, Häusser M. 2019. Predictive and reactive reward signals conveyed by climbing fiber inputs to cerebellar Purkinje cells. Nat Neurosci 22: 950–962.

Liang B, Zhang L, Barbera G, Fang W, Zhang J, Chen X, Chen R, Li Y, Lin D-T. 2018. Distinct and Dynamic ON and OFF Neural Ensembles in the Prefrontal Cortex Code Social Exploration. Neuron 100: 700–714.e9.

Liberti WA 3rd, Markowitz JE, Perkins LN, Liberti DC, Leman DP, Guitchounts G, Velho T, Kotton DN, Lois C, Gardner TJ. 2016. Unstable neurons underlie a stable learned behavior. Nat Neurosci 19: 1665–1671.

Liberti WA, Perkins LN, Leman DP, Gardner TJ. 2017. An open source, wireless capable miniature microscope system. J Neural Eng 14: 045001.

Mallet N, Le Moine C, Charpier S, Gonon F. 2005. Feedforward inhibition of projection neurons by fast-spiking GABA interneurons in the rat striatum in vivo. J Neurosci 25: 3857–3869.

Murugan M, Jang HJ, Park M, Miller EM, Cox J, Taliaferro JP, Parker NF, Bhave V, Hur H, Liang Y, Nectow AR, Pillow JW, Witten IB. 2017. Combined Social and Spatial Coding in a Descending Projection from the Prefrontal Cortex. Cell 171: 1663–1677.e16.

Oñativia J, Schultz SR, Dragotti PL. 2013. A finite rate of innovation algorithm for fast and accurate spike detection from two-photon calcium imaging. J Neural Eng 10: 046017.

Pnevmatikakis EA, Giovannucci A. 2017. NoRMCorre: An online algorithm for piecewise rigid motion correction of calcium imaging data. J Neurosci Methods 291: 83–94.

Prudente CN, Stilla R, Buetefisch CM, Singh S, Hess EJ, Hu X, Sathian K, Jinnah HA. 2015. Neural Substrates for Head Movements in Humans: A Functional Magnetic Resonance Imaging Study. J Neurosci 35: 9163–9172.

Quy PN, Fujita H, Sakamoto Y, Na J, Sugihara I. 2011. Projection patterns of single mossy fiber axons originating from the dorsal column nuclei mapped on the aldolase C compartments in the rat cerebellar cortex. J Comp Neurol 519: 874–899.

R Core Team. 2019. R: A Language and Environment for Statistical Computing.

Remedios R, Kennedy A, Zelikowsky M, Grewe BF, Schnitzer MJ, Anderson DJ. 2017. Social behaviour shapes hypothalamic neural ensemble representations of conspecific sex. Nature 550: 388–392.

Romano V, De Propris L, Bosman LW, Warnaar P, Ten Brinke MM, Lindeman S, Ju C, Velauthapillai A, Spanke JK, Middendorp Guerra E, Hoogland TM, Negrello M, D’Angelo E, De Zeeuw CI. 2018. Potentiation of cerebellar Purkinje cells facilitates whisker reflex adaptation through increased simple spike activity. Elife 7. doi: 10.7554/eLife.38852

Ruigrok TJH. 2010. Ins and Outs of Cerebellar Modules. Cerebellum 464–474.

Sauerbrei B, Guo J-Z, Mischiati M, Guo W, Kabra M, Verma N, Branson K, Hantman A. 2018. Motor cortex is an input-driven dynamical system controlling dexterous movement. bioRxiv. doi: 10.1101/266320

Schindelin J, Arganda-Carreras I, Frise E, Kaynig V, Longair M, Pietzsch T, Preibisch S, Rueden C, Saalfeld S, Schmid B, Tinevez J-Y, White DJ, Hartenstein V, Eliceiri K, Tomancak P, Cardona A. 2012. Fiji: an open-source platform for biological-image analysis. Nat Methods 9: 676–682.

Stamatakis AM, Schachter MJ, Gulati S, Zitelli KT, Malanowski S, Tajik A, Fritz C, Trulson M, Otte SL. 2018. Simultaneous Optogenetics and Cellular Resolution Calcium Imaging During Active Behavior Using a Miniaturized Microscope. Front Neurosci 12: 496.

Stirman JN, Smith IT, Kudenov MW, Smith SL. 2016. Wide field-of-view, multi-region, two-photon imaging of neuronal activity in the mammalian brain. Nat Biotechnol 34: 857–862.

Stoodley CJ, D’Mello AM, Ellegood J, Jakkamsetti V, Liu P, Nebel MB, Gibson JM, Kelly E, Meng F, Cano CA, Pascual JM, Mostofsky SH, Lerch JP, Tsai PT. 2017. Altered cerebellar connectivity in autism and cerebellar-mediated rescue of autism-related behaviors in mice. Nat Neurosci 20: 1744–1751.

Tecuapetla F, Matias S, Dugue GP, Mainen ZF, Costa RM. 2014. Balanced activity in basal ganglia projection pathways is critical for contraversive movements. Nat Commun 5: 4315.

Terada S-I, Kobayashi K, Ohkura M, Nakai J, Matsuzaki M. 2018. Super-wide-field two-photon imaging with a micro-optical device moving in post-objective space. Nat Commun 9: 3550.

van Rossum G. 1995. Python tutorial. Amsterdam: Centrum voor Wiskunde en Informatica (CWI).

Wagner MJ, Kim TH, Kadmon J, Nguyen ND, Ganguli S, Schnitzer MJ, Luo L. 2019. Shared Cortex-Cerebellum Dynamics in the Execution and Learning of a Motor Task. Cell 0. doi: 10.1016/j.cell.2019.02.019

Wagner MJ, Kim TH, Savall J, Schnitzer MJ, Luo L. 2017. Cerebellar granule cells encode the expectation of reward. Nature 544: 96–100.

Witter L, Canto CB, Hoogland TM, de Gruijl JR, De Zeeuw CI. 2013. Strength and timing of motor responses mediated by rebound firing in the cerebellar nuclei after Purkinje cell activation. Front Neural Circuits 7: 133.

Zhou P, Resendez SL, Rodriguez-Romaguera J, Jimenez JC, Neufeld SQ, Giovannucci A, Friedrich J, Pnevmatikakis EA, Stuber GD, Hen R, Kheirbek MA, Sabatini BL, Kass RE, Paninski L. 2018. Efficient and accurate extraction of in vivo calcium signals from microendoscopic video data. Elife 7. doi: 10.7554/eLife.28728

